# Adult oligodendrogenesis gates arcuate neuronal glucose sensing through remodelling of the blood-hypothalamus barrier via ADAMTS4

**DOI:** 10.1101/2024.09.10.612227

**Authors:** Sophie Buller, Emily O. Staricoff, Christine Riches, Anthony Tsang, Masa Josipovic, Kentaro Ikemura, Gabriel Opoku, Ikumi Sato, Satoshi Hirohata, Saskia Stenzel, Stuart G. Nayar, Marta Ramos Vega, Jacob Hecksher-Sørensen, Sebastian Timmler, Georgina K.C. Dowsett, Brian Y.H. Lam, Giles S.H. Yeo, Kimberly M. Alonge, Huiliang Li, William D. Richardson, Mark L. Evans, Clemence Blouet

## Abstract

Brain glucose sensing is critical for survival during hypoglycaemia and tunes the level of defended blood glucose, which goes up in diabetes. Neuronal glucose sensing neurons and mechanisms have been identified, but how these neurons access blood concentrations of glucose to adjust their output and maintain glucose homeostasis is unclear. Here, we demonstrate that adult oligodendrogenesis in the median eminence (ME) is modulated by changes in circulating glucose levels and rapidly upregulated by hypoglycaemia. We show that genetic blockade of new OL production in adult mice impairs the regulation of glucose homeostasis, the integrity of the ME blood-hypothalamus barrier, and neuronal glucose sensing in the arcuate nucleus of the hypothalamus (ARH). Unexpectedly, functional integrity of adult-formed myelin is not required for the maintenance of glucose homeostasis. Instead, newly formed OLs exert their glucoregulatory actions via the synthesis of A disintegrin and metallopeptidase with thrombospondin motifs 4 (ADAMTS4), a metallopeptidase expressed exclusively by OLs and dependent on adult OL genesis to maintain its expression in the ME. Both lack of *Adamts4* and ADAMTS4 gain-of-function are associated with impaired glucose homeostasis and remodelling of the blood-hypothalamus barrier, indicating that optimal ADAMTS4 expression is required for the integrity of vascular permeability and normal glycaemic control. Finally, we show that ME ADAMTS4 expression is regulated by changes in peripheral glycaemia and is dysregulated in diabetes, providing a mechanism by which ME OLs contribute to the regulation of glucose homeostasis.

## Main

The brain plays a critical role in the regulation of glucose homeostasis, not only by regulating pancreatic insulin release and peripheral insulin action^1^, but also through its key role in mounting the neuroendocrine counterregulatory response to hypoglycaemia^2^. This essential survival response is coordinated by neurocircuitries distributed between the hindbrain and mediobasal hypothalamus (MBH), enriched in hypoglycaemia-activated neurons. Glucose sensing neurons of the ventromedial hypothalamic nucleus (VMN) are required for the production of the counterregultory response to hypoglycaemia. How these neurons access circulating concentrations of glucose is unclear, but might rely on their proximity to the median eminence (ME). Here the fenestrated vasculature exposes nearby neurons to unbuffered concentrations of circulating glucose, a feature which is likely critical for the production of rapid and accurate neuroendocrine responses mediating the return to homeostasis^3–5^.

Located adjacent to the arcuate nucleus of the hypothalamus (ARH), the ME is highly plastic under physiological conditions and rapidly remodelled in response to hypoglycaemia, leading to increased blood-brain-barrier (BBB) permeability and/or blood-parenchymal diffusion and greater access of circulating signals to adjacent nuclei^6^. These modifications are largely attributed to β_2_-tanycytes, specialised ependymal cells that line the 3^rd^ ventricle and exhibit glucose-sensing properties^7–9^, yet a role for other cell types in hypoglycaemia-induced remodelling of the ME is unexplored.

Emerging evidence indicates that oligodendrocyte (OL) lineage cells display high levels of plasticity in the ME during adulthood^10,11^. ME oligodendrogenesis continues in the healthy adult brain at high rate and is regulated by nutritional signals^10,11^, yet the specific cue(s) that regulate oligodendrocyte progenitor cell (OPC) differentiation are unknown. Although the functional relevance of ME OL plasticity is yet to be fully elucidated, available data suggests that new OL generation in adulthood benefits metabolic regulation and promotes hypothalamic leptin sensitivity^10^. Given that MBH neuroglial interactions are important determinants of systemic glucose homeostasis with roles for astrocytes^12,13^, tanycytes^14^ and microglia^15,16^ described, here we explored the hypothesis that ME OL plasticity is regulated by peripheral blood glucose levels and contributes to the regulation of glucose homeostasis in adult mice.

## Results

### Acute changes in glycaemia regulate oligodendrocyte differentiation in the adult median eminence

To determine how changes in blood glucose levels regulate ME oligodendrocyte lineage cells, we first characterised ME OPC proliferation in response to acute changes in blood glucose levels. C57BL/6J mice received intraperitoneal (ip) injections of vehicle (saline), glucose, 2-deoxy-d-glucose (2DG, a non-metabolizable glucose analogue) or insulin, one hour prior to perfusion-fixation. As expected, glucose and 2DG administration resulted in hyperglycaemia (the latter reflecting the counter-regulatory response to glucoprivation) and insulin led to hypoglycaemia (**Extended Data Fig. 1a**). All mice also received bromodeoxyuridine (BrdU), a thymidine analogue that is incorporated into the DNA of actively dividing cells, to label proliferating cells as previously described^17^. Immunostaining against pan-OL marker SRY-box transcription factor 10 (SOX10) and OPC marker platelet derived growth factor receptor alpha (PDGFRα) was used to identify OPCs (SOX10^+^/PDGFRα^+^, **Fig. 1a)**. Remarkably within 1h, 2DG significantly increased OPC proliferation (SOX10^+^/PDGFRα^+^/BrdU^+^) in the ME, leading to an increase in OPC density (**Fig. 1b-d**). Glucose treatment resulted in a near-significant increase in OPC proliferation (SOX10^+^/PDGFRα^+^/BrdU^+^; **Fig. 1d**). Thus, hyperglycaemia rapidly promotes ME OPC proliferation. In contrast, insulin treatment did not affect OPC proliferation or density (**Fig. 1b-d**).

**Figure 1:**
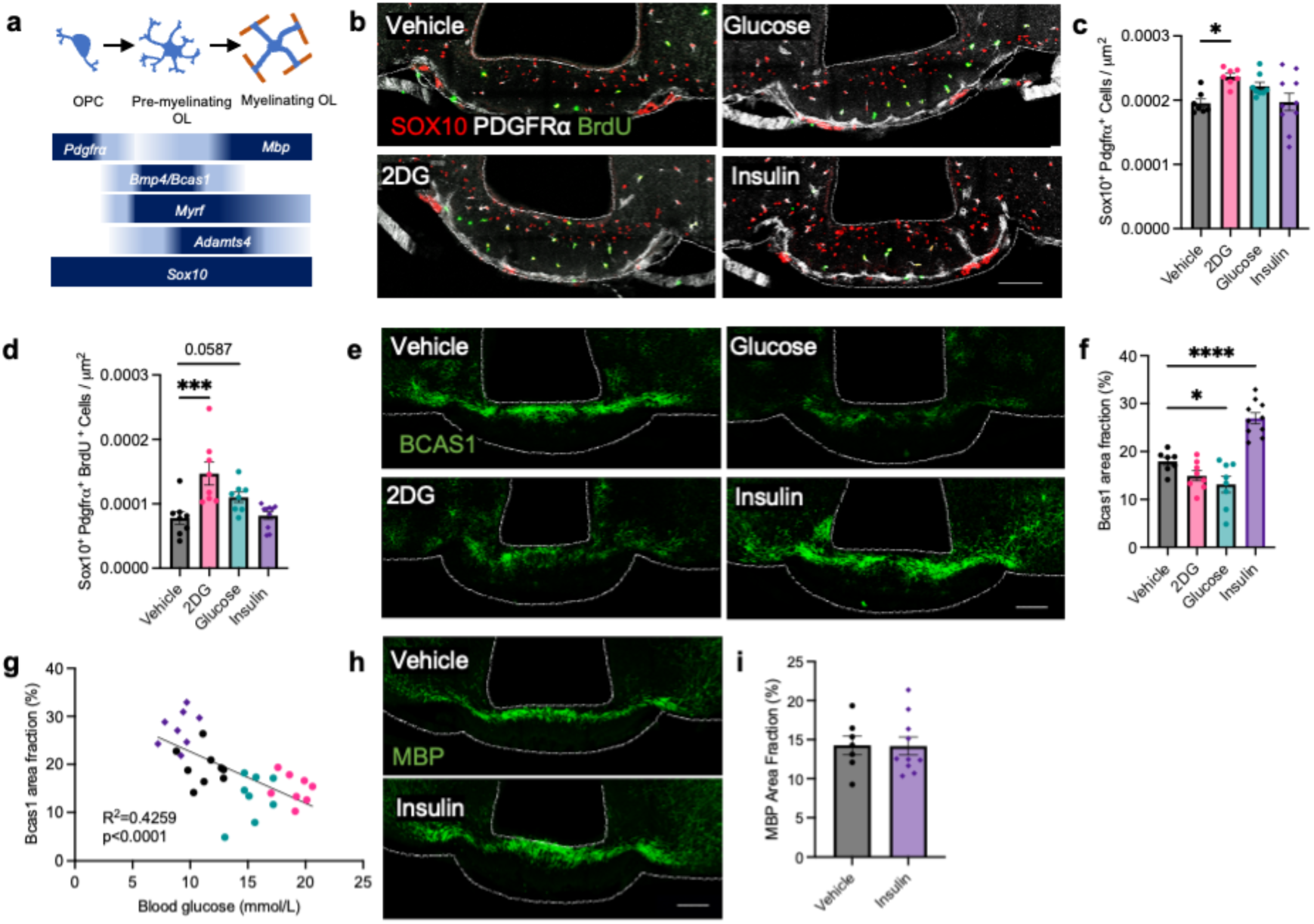
Acute changes in glycaemia regulate oligodendrocyte differentiation in the median eminence. (**a**) Markers of OL lineage cell stages in the adult brain. **(b**) Representative images of pan-OL lineage marker SOX10, OPC marker PDGFRa and Bromodeoxyuridine (BrdU) in the mediobasal hypothalamus (MBH) of C57BL/6J mice one hour after the intraperitoneal administration of vehicle (saline; 10 ml/kg), 2-deoxy-d-glucose (2DG; 250 mg/kg), glucose (2 g/kg) or insulin (0.75 U/kg) and the quantification of (**c**) OPCs (SOX10+/PDGFRa+) and (**d**) proliferating (BrdU+) OPCs in the ME. (**e**) Representative images of Breast carcinoma amplified sequence 1 (BCAS1), a marker of newly differentiated, pre-myelinating OLs in the MBH following vehicle, 2DG, glucose or insulin administration, (**f**) the quantification of BCAS1 expression in the ME and (**g**) its correlation with blood glucose. (**h**) Representative images of Myelin basic protein (MBP), a myelin structural protein, in the MBH of mice that received vehicle or insulin and (**i**) associated quantification. All data presented as mean ± S.E.M., scale bars represent 100 μm. Data analysed using one-way ANOVA or Kruskal-Wallis test with Dunnett’s- or Dunn’s-multiple comparisons test when comparing 4 independent groups or an unpaired student’s t test when comparing 2 independent groups, *p<0.05, **p<0.01, ***p<0.001, ****p<0.0001. Correlations were analysed using simple linear regression, n=8-10/group.

Next, to assess whether changes in glycaemia regulate OPC differentiation into OLs in the ME, we analysed the expression of breast carcinoma amplified sequence 1 (BCAS1), a marker of newly formed, pre-myelinating OLs^18^. 2DG administration had no effect on ME BCAS1 immunoreactivity. In contrast, BCAS1 expression significantly decreased following glucose administration and robustly increased in response to insulin administration (**Fig. 1e-f**), suggesting that deviations from euglycaemia bidirectionally regulate ME oligodendrogenesis. In fact, blood glucose levels inversely correlate with ME BCAS1 expression (**Fig. 1g**). Of note, despite the lack of change in OPC density and proliferation following insulin treatment, the density of OL lineage cells (SOX10^+^; representing OPCs and OLs) significantly increased (**Extended Data Fig. 1b**) suggesting that insulin increases OL survival in the ME. However, expression of myelin basic protein (MBP) remained unchanged in the ME between vehicle and insulin treated groups (**Fig. 1h-i**).

In insulin treated animals, either hyperinsulinemia or subsequent hypoglycaemia could drive the response of ME OL lineage cells. To tease these effects apart, we performed hyperinsulinaemic hypoglycaemic, euglycaemic and hyperglycaemic clamps in conscious-free moving mice (**Fig. 2a-c**). Perfusion-fixed tissues were collected after 75-minutes for analysis of Bone morphogenetic protein 4 (*Bmp4)*, a specific marker of newly differentiated OLs^19^, by single molecule fluorescent *in situ* hybridisation (sm-FISH, RNAscope technology; **Fig. 2d**). Consistent with our results indicating increased OPC differentiation into new OLs following insulin administration, *Bmp4* expression was significantly increased in the ME of mice following a 75-minute hyperinsulinaemic hypoglycaemic clamp (**Fig. 2e-f**) confirming that hypoglycaemia rapidly increases oligodendrogenesis in the ME.

**Figure 2:**
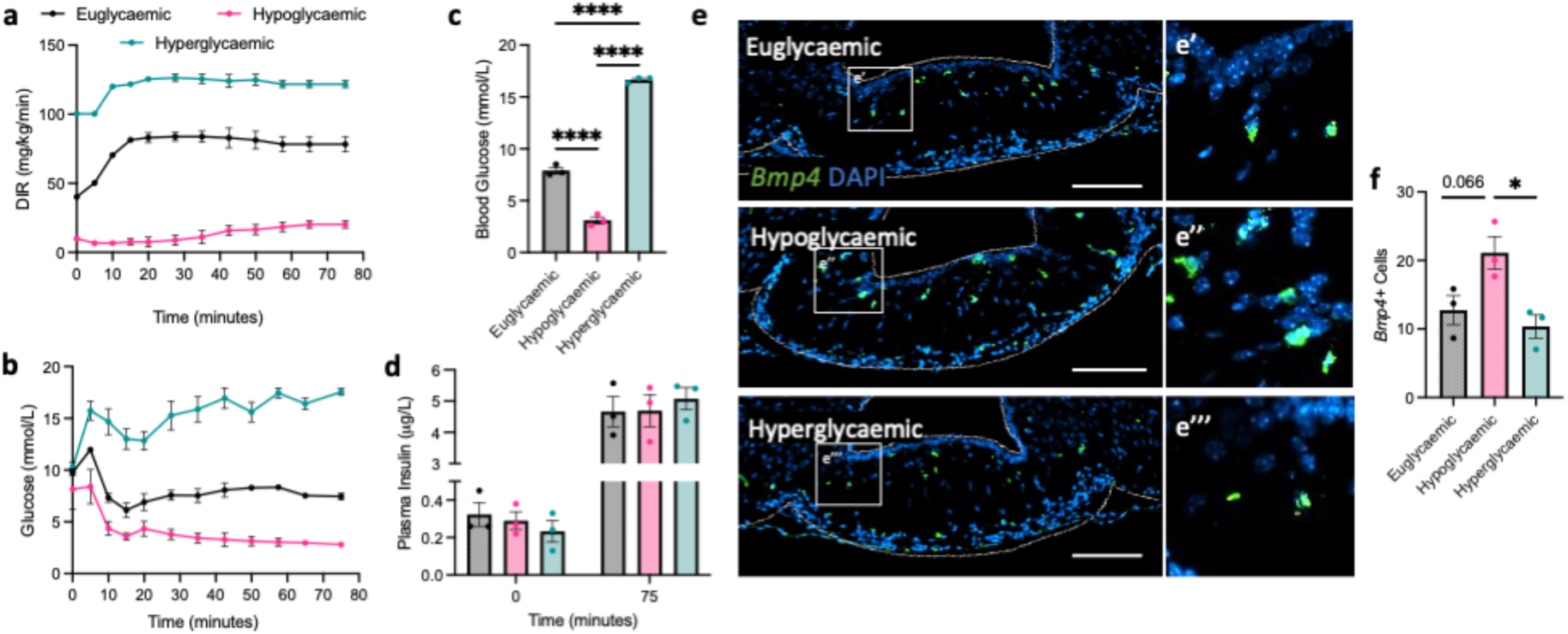
Hypoglycaemia rapidly increases median eminence oligodendrocyte progenitor cell differentiation. (**a**) Dextrose infusion rates (DIR; mg/kg/min) and (**b**) blood glucose levels (mmol/L) during a 75-minute hyperinsulinaemic-euglycaemic, - hypoglycaemic or hyperglycaemic clamp, (**c**) average blood glucose and (**d**) plasma insulin levels at 0 and 75 minutes. (**e**) Representative images of RNAscope single-molecule fluorescence *in situ* hybridization in the median eminence of C57BL/6J mice exposed to a 75-minute hyperinsulinaemic-euglycaemic, hypoglycaemic or hyperglycaemic clamp. Probes were used to label *Bone morphogenetic protein 4* (*Bmp4*) – expressed by newly differentiated OLs. (**f**) Quantification of the number of cells expressing each marker, data presented as mean ± S.E.M. and analysed using one-way ANOVA with Tukey’s multiple comparisons test *p<0.05, n=3/group, scale bars represent 100 μm.

### Adult oligodendrogenesis contributes to the counterregulatory response to hypoglycaemia and the maintenance of glucose homeostasis

Glucose sensing in the MBH initiates the neuroendocrine response to hypoglycaemia which restores systemic glucose homeostasis^6^. To determine whether increased ME OPC differentiation into new OLs during hypoglycaemia contributes to this function, we used *Pdgfra-CreER^T2^;Rosa26R-eYFP;Myrf^fl/fl^* (*Myrf^fl/fl^*) mice, where conditional deletion of myelin regulatory factor (*Myrf*) in OPCs blunts new OL production following tamoxifen administration at postnatal day 60 (P60)^10^, and examined their glycaemic control. Compared to wild-type littermates (*Myrf^+/+^*), *Myrf^fl/fl^* mice became resistant to the hypoglycaemic action of insulin within six weeks of tamoxifen administration (**Fig. 3a-b**). Consistent with the development of insulin resistance, plasma insulin levels in both the fasted and *ad libitum* fed state were significantly increased in *Myrf^fl/fl^* mice (**Fig. 3c**). When challenged with 2DG, *Myrf^fl/fl^* mice mounted an exacerbated hyperglycaemic (**Fig. 3d-e, Extended Data Fig. 2a-b**) and hyperphagic (**Fig. 3f, Extended Data Fig. 2c**) response, indicating an impaired counterregulatory response to neuroglycopenia. Finally, *Myrf^fl/fl^* mice displayed glucose intolerance when challenged with an oral gavage of glucose (**Fig. 3g-h**). However, glucose-induced hyperinsulinemia did not differ between groups (**Fig. 3i**) nor did plasma glucagon levels in unchallenged mice (**Fig. j**) suggesting normal pancreatic function following *Myrf* deletion. Collectively these data support a role for adult born OLs in the response to glucose deficit and the activation of the counterregulatory response to hypoglycaemia.

**Figure 3:**
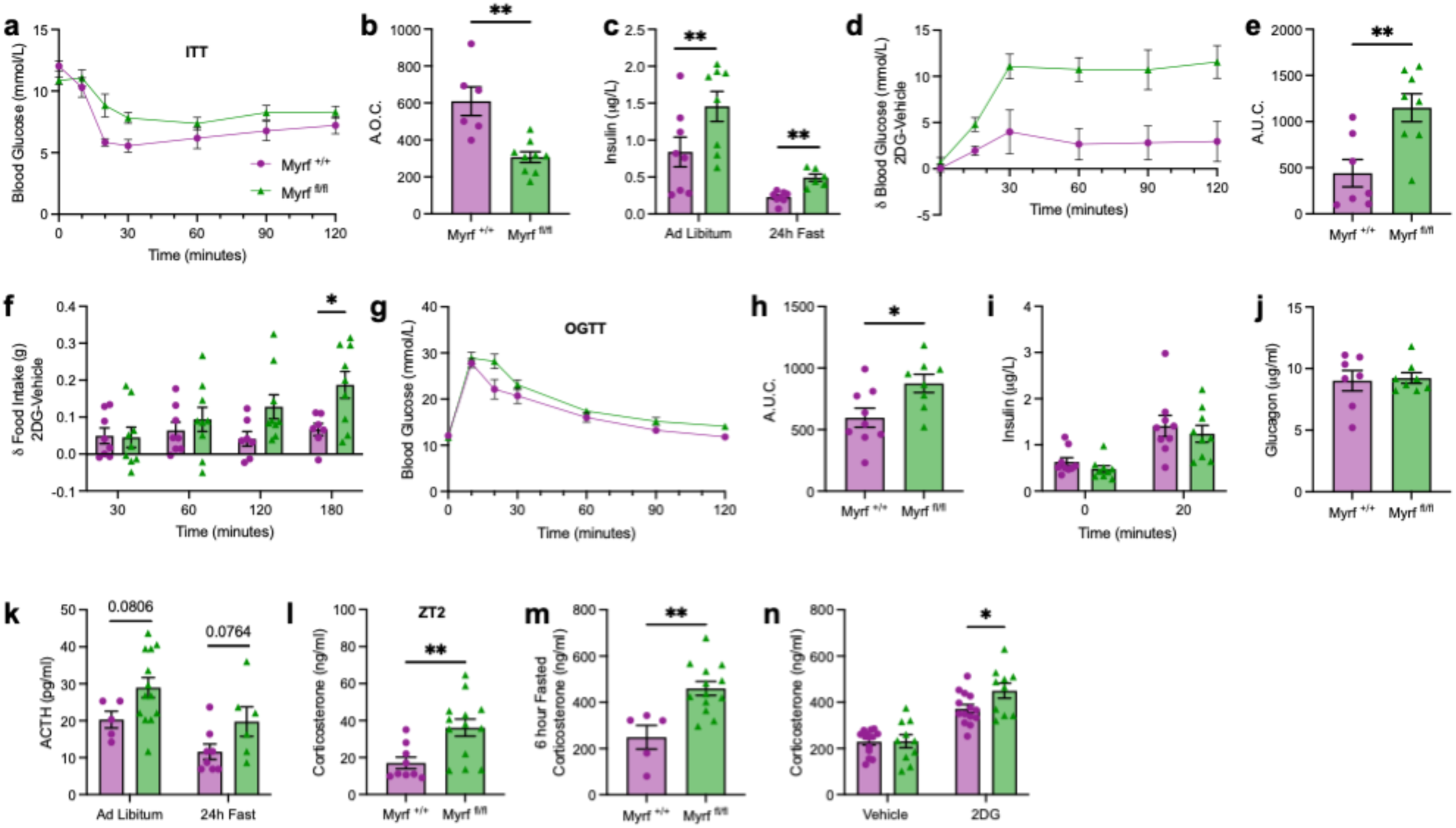
Adult oligodendrocyte differentiation is required for glucose homeostasis. (**a**) Blood glucose levels in *Myrf ^­­^*and *Myrf ^­­^*mice over 120 minutes following an intraperitoneal injection of insulin (0.75 U/kg) and (**b**) the resulting area over the curve. (**c**) Plasma insulin levels in *ad libitum* fed and 24 hour fasted *Myrf ^­­^*and *Myrf ^­­^*mice. (**d**) Blood glucose levels in *Myrf ^­­^*and *Myrf ^­­^*mice over 120 minutes following an intraperitoneal injection of 2-deoxy-d-glucose (2DG; 250 mg/kg) compared to vehicle (saline; 10 ml/kg) and (**e**) the resulting area under the curve. (**f**) Food intake over 180 minute in *Myrf ^­­^*and *Myrf ^­­^*mice following an intraperitoneal injection of 2DG compared to vehicle. (**g**) Blood glucose levels in *Myrf ^­­^*and *Myrf ^­­^*mice over 120 minutes after an oral gavage of glucose (2 g/kg), (**h**) the resulting area under the curve and (**i**) plasma insulin levels at baseline and after 20 minutes. (**j**) Plasma glucagon levels in unchallenged *Myrf ^­­^*and *Myrf ^­­^*mice. (**k)** Plasma adrenocorticotrophic hormone levels in *ad libitum* fed and 24 hour fasted *Myrf ^­­^*and *Myrf ^­­^*mice. Plasma corticosterone levels in *Myrf ^­­^*and *Myrf ^­­^*mice at (**l**) zeitgeber time (ZT) 2 and (**m**) after a 6 hour fast. (**n**) Plasma corticosterone levels in *Myrf ^­­^*and *Myrf ^­­^*mice 30 minutes after vehicle or 2DG administration. All data presented as mean ± S.E.M.. Data analysed using an unpaired student’s test or two-way ANOVA with Sidak’s multiple comparisons test *p<0.05, **p<0.01, n=6-14/group.

The glucose counterregulatory response which orchestrates the recovery from hypoglycaemia involves activation of the hypothalamic pituitary adrenal (HPA) axis through the release of corticotropin-releasing hormone (CRH) in the ME from the hypothalamic paraventricular nucleus (PVH). CRH reaches the anterior pituitary through the hypophyseal portal circulation to stimulate ACTH synthesis, which in turns stimulates corticosterone secretion from the adrenal cortex. Thus, we next examined the activity of the HPA axis during blockade of adult oligodendrogenesis. In *Myrf^fl/fl^* mice, plasma levels of adrenocorticotrophic hormone (ACTH) were increased to near statistical significance under *ad libitum* feeding conditions at zeitgeber time (ZT) 2 or following a 24 hour fast (**Fig. 3k**). Consistently, plasma corticosterone levels were significantly elevated in *Myrf^fl/fl^* mice at ZT2 (**Fig. 3l**) and after a 6 hour fast (**Fig. 3m**), indicating increased HPA axis activity. Finally, we assessed the ability of *Myrf^fl/fl^*mice to mount an appropriate neuroendocrine response to glucoprivation, which involves increases in circulating levels of corticosterone and glucagon to prevent a further decline in plasma glucose levels and restore euglycaemia^2^. Consistent with a heightened sensitivity to glucoprivation, *Myrf^fl/fl^* mice displayed an exacerbated hypercorticosteronaemic, but not hyperglucagonemic response to 2DG (**Fig. 3n, Extended Data Fig. 2d**). Collectively, these data support a role for adult oligodendrogenesis in the regulation of the HPA axis and the activation of counterregulatory mechanisms in response to hypoglycaemia.

### Adult oligodendrogenesis regulates blood-hypothalamus barrier plasticity and ARH glucose sensing

During hypoglycaemia, the blood-hypothalamus barrier is reorganised to increase access of circulating signals to ARH neurocircuits, at least in part through modification of tanycytic tight junction complexes^6^. However, whether OL plasticity contributes to these adaptations remains unexplored. Therefore, we first asked whether blunted OL differentiation in *Myrf^fl/fl^* mice impacts MBH tanycytes. ME-ARH tanycytic morphology and density were unchanged in *Myrf^fl/fl^* mice (**Extended Data Fig. 3a-b**), however the expression of the tight-junction protein Zona occludens 1 (ZO-1) was increased in both the dorsal and ventral ME (**Fig. 4a-b**), suggesting decreased paracellular permeability and remodelling of local barriers. The number of MECA32+/Laminin+ capillary loops was increased in the ME of *Myrf^fl/fl^* mice (**Fig. 4c-d**) indicating increased blood vessel fenestration and BHB permeability. Consistently, Evans Blue infiltration into the ME-ARH following an intra-vascular bolus was greater in *Myrf^fl/fl^* mice compared to controls (**Fig. 4e-f**), indicating impaired ME-ARH barrier function and increased access of peripheral signals to the hypothalamus. Thus, the structural organisation of the ME in *Myrf^fl/fl^* mice resembles functional adaptations of the ME that occur during hypoglycaemia^6^, suggesting that OL differentiation contributes to these adaptations and influence ARH glucose sensing. To address this question, we assessed cFOS expression in the ARH of *Myrf^fl/fl^*mice and littermate controls 90 minutes after ip administration of saline or 2DG (**Fig. 4g**). In both *Myrf^+/+^* and *Myrf^fl/fl^* mice, 2DG treatment induced robust cFOS expression in the ARH compared to vehicle. However, consistent with the exacerbated hyperphagic and hyperglycaemic response to 2DG in *Myrf^fl/fl^* mice, the number of cFOS^+^ cells in the ARH was significantly higher in *Myrf^fl/fl^* mice compared to controls following treatment with 2DG (**Fig. 4h**). Thus, OPC differentiation into new OLs regulates hypothalamic neuronal glucose sensing.

**Figure 4:**
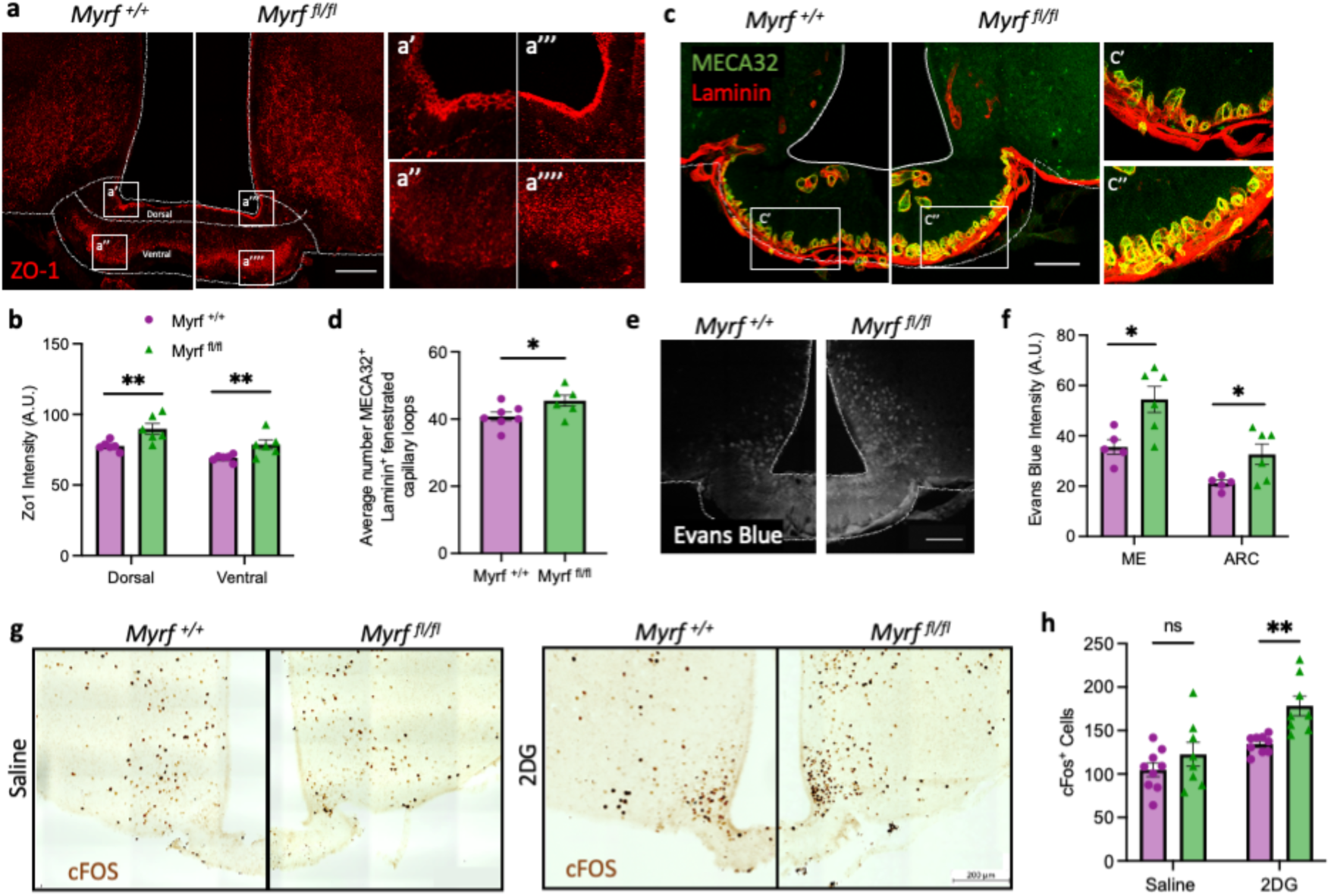
Adult oligodendrocyte differentiation regulates glucose sensing in the mediobasal hypothalamus. (**a**) Representative images of Zona occludens 1 (ZO-1) immunolabelling in the mediobasal hypothalamus (MBH) of myelin regulatory factor (MYRF) conditional knockout mice (*Myrf^­­^*) and controls (*Myrf^­­^*) and (**b**) associated quantification. (**c**) Representative images of laminin and pan-endothelial cell marker MECA-32 in the MBH of *Myrf^­­^*and *Myrf^­­^*mice and (**d**) associated quantification. (**e**) Evans blue permeation of the mediobasal hypothalamus (MBH) in *Myrf^­­^*and *Myrf^­­^*mice following an intravascular bolus at zeitgeber time 2 and (**f**) associated quantification. (**g**) Representative images of cFOS immunohistochemistry in the MBH 90 minutes after an intraperitoneal injection of saline (10 ml/kg) or 2-deoxy-d-glucose (2DG; 250 mg/kg) to *Myrf ^­­^*and *Myrf ^­­^*mice and (**h**) associated quantification of cFOS+ cells in the arcuate nucleus (ARC). Data presented as mean ± S.E.M., scale bars represent 100 μm. Data analysed using either an unpaired student’s- or Welch’s-t-test, or 2-way ANOVA with Sidak’s multiple comparisons test, *p<0.05, **p<0.01, n=5-9/group.

### New compact myelin generation is not required for the regulation of glucose homeostasis

In the ME, myelin rapidly turns over such that genetic block of new OL differentiation in adult *Myrf^fl/fl^* mice leads to rapid and significant reduction in ME MBP expression^10^. Since myelin ensheaths axons of magnocellular arginine vasopressin (AVP) neurones (**Fig. 5a**), which pass though the ME and have been implicated in the regulation of glucose homeostasis^20^, demyelination of ME axons in *Myrf^fl/fl^* mice could mediate their neuroendocrine and glycaemic phenotype. To test this hypothesis, we used *Pdgfra-CreER^T2^;Rosa26R-eYFP;Mbp^fl/fl^*(*Mbp^fl/fl^*) mice in which tamoxifen-induced deletion of the *Mbp* gene promoter from OPCs blocks *Mbp* transcription, MBP protein accumulation and myelin compaction in newly-formed myelin sheaths (SGN, HL and WDR, unpublished). This allows us to specifically assess the contribution of adult-formed myelin to the regulation of glucose homeostasis. Deletion of *Mbp* in OPCs led to a robust (∼50%) reduction in ME MBP expression 6 weeks after tamoxifen administration at P60 in *Mbp^fl/fl^* mice (**Fig. 5b-c**) and was associated with a decrease in the total number of OL lineage cells (SOX10+) and a near-significant reduction in OL (Sox10^+^/PDGFRα^-^) density in the ME (**Extended Data Figure 4a-b**). We then used serial electron microscopy (SEM) to assess myelin ultrastructure in the ME of both *Myrf^fl/fl^* and *Mbp^fl/fl^* mice compared to wild-type controls (**Fig. 5d**). As expected, both deletion of *Myrf* and *Mbp* from OPCs significantly reduced the number of myelinated axons in the ME (**Fig. 5e**) and quantification of the g-ratio on axons displaying compact myelin sheaths demonstrated a significant increase in the g-ratio in both *Myrf^fl/fl^* and *Mbp^fl/fl^* mice compared to wild-type controls indicating thinner myelin sheathes (**Fig. 5f-g**). Despite changes in ME myelin density and structure and the number of cells in the OL lineage, the density of newly formed OLs (Sox10^+^/PDGFRα^-^/YFP^+^; **Extended Data Fig. 4a-c**) did not change in the ME following conditional deletion of *Mbp* in OPCs. Consistently, BCAS1 expression was comparable between *Mbp^fl/fl^* mice and littermate controls (**Extended Data Fig. 4d-e**) indicating that deletion of *Mbp* from adult OPCs does not impact ME OPC differentiation or new OL production, and specifically impairs compactness of adult-formed myelin.

**Figure 5:**
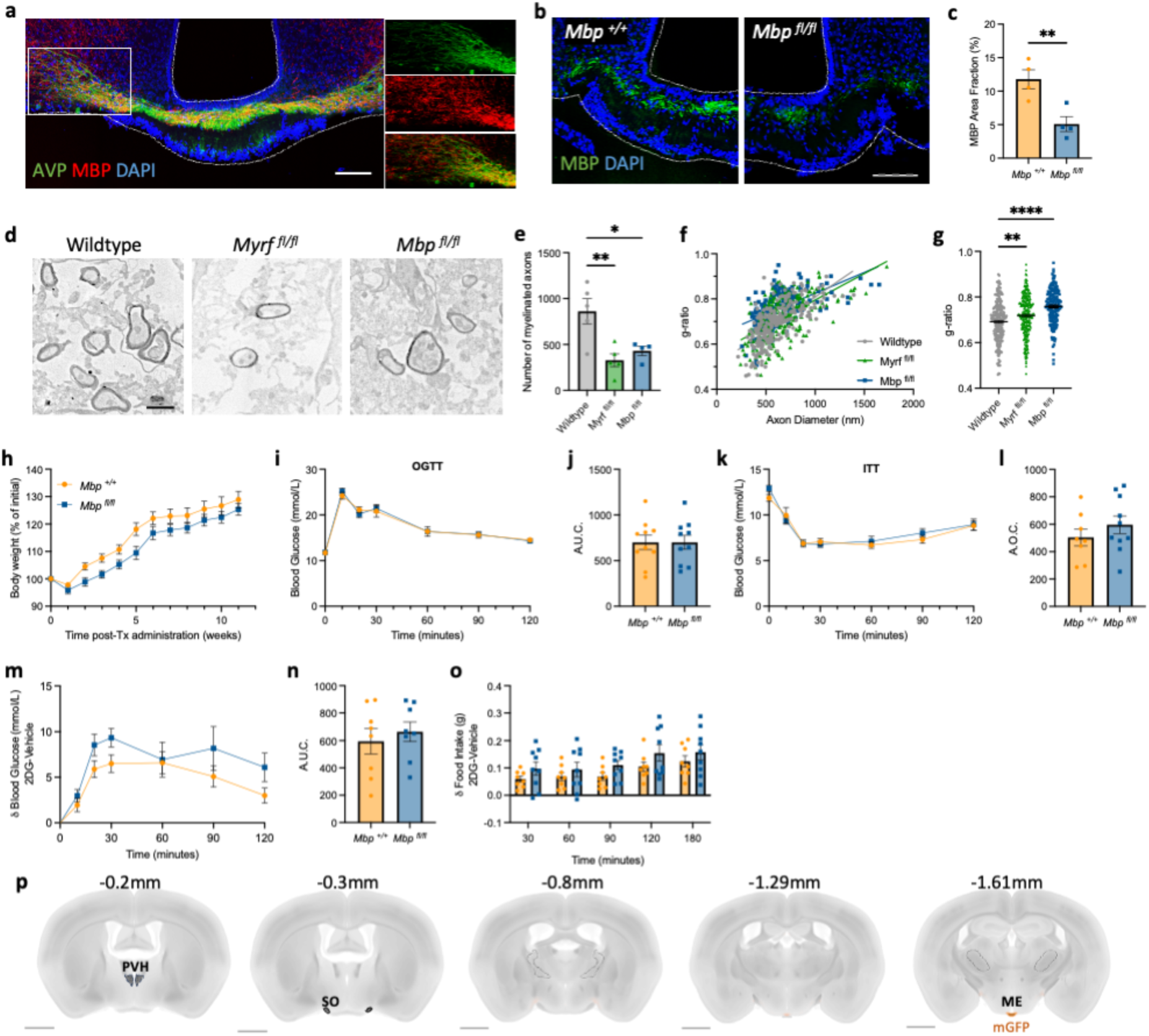
Adult myelin plasticity does not regulate glucose homeostasis in adult mice. (**a**) Representative images demonstrating myelination of arginine vasopressin (AVP) axons in the mediobasal hypothalamus. (**b**) Representative images of myelin basic protein (MBP) immunolabelling in the median eminence (ME) of MBP conditional knockout mice (*Mbp ^­­^*) and controls (*Mbp ^­­^*) six weeks after tamoxifen administration at postnatal day 60 and (**c**) associated quantification. (**d**) Representative images of myelin ultrastructure and integrity in the ME of wildtype, myelin regulatory factor conditional knockout (*Myrf ^­­^) and Mbp ^­­^*mice by serial transmission electron microscopy and associated quantifications of (**e**) the number of myelinated axons and (**f-g**) g ratio. (**h**) Body weight change over 12 weeks after tamoxifen administration to *Mbp ^­­^*and *Mbp ^­­^*mice. (**i**) Blood glucose over 120 minutes after an oral gavage of glucose (2 g/kg) in *Mbp ^­­^*and *Mbp ^­­^*mice and (**j**) the associated area under the curve. (**k**) Blood glucose over 120 minutes following an intraperitoneal injection of insulin (0.75 U/kg) and (**l**) the associated area over the curve. (**m**) Blood glucose levels in *Mbp ^­­^*and *Mbp ^­­^*mice over 120 minutes following an intraperitoneal injection of 2-deoxy-d-glucose (2DG; 250 mg/kg) compared to vehicle (saline; 10 ml/kg) and (**n**) the resulting area under the curve. (**o**) Food intake over 180 minute in *Mbp ^­­^*and *Mbp ^­­^*mice following an intraperitoneal injection of 2DG compared to vehicle. (**p**) Coronal maps of mGFP fluorescence in *Pdgfrα-CreERT2;taumGFP* mice dosed with tamoxifen at P60 and perfused at P90. mGFP is specifically expressed in the ME, demonstrating that only the terminal portion of magnocellular axons within the ME are exposed to myelin plasticity. PVH: paraventricular nucleus of the hypothalamus, SO: supraoptic nucleus. Scale bar is 1 mm. All data presented as mean ± S.E.M., scale bars represent 100 μm. Data analysed using an unpaired student’s test, one-way ANOVA with Dunnett’s multiple comparisons test or two-way ANOVA with Sidak’s multiple comparisons test, *p<0.05, **p<0.01, ****p<0.0001 n=4-9/group.

Phenotypically, deletion of *Mbp* in OPCs at P60 produced a trend towards reduced weight gain (genotype effect p=0.0803, interaction (time*genotype) p=0.0725; **Fig. 5h**). However, unlike what we observed in *Myrf^fl/fl^* mice, glucose tolerance, insulin sensitivity and the response to 2DG-induced glucoprivation did not differ between *Mbp^fl/fl^* mice and wild-type littermates (**Fig. 5i-o, Extended Data Fig. 4f-h**). Thus, the adult generation of new compact myelin is dispensable for the regulation of glucose homeostasis in adult mice.

To further characterise the functional consequences of ME demyelination in *Myrf^fl/fl^* mice, we assessed the integrity of the AVP axis in *Myrf^fl/fl^* mice focussing on their role in osmoregulation. ME AVP immunolabelling was unchanged in *Myrf^fl/fl^* mice (**Extended Data Fig. 4i-j**) and *ad libitum* water intake was comparable to controls **(Extended Data Fig. 4k**). Furthermore, *Myrf^fl/fl^* mice exhibited the expected increase in plasma osmolarity during a dehydration challenge (**Extended Data Fig. 4l**), suggesting intact osmoregulation. Thus, consistent with the phenotype of *Mbp^fl/fl^*mice, these data indicate that new adult OL and/or myelin production is not required for the integrity of the vasopressin axis. Extending on these finding, we visualised new myelin production in the adult brain between P60 and P90 in *Pdgfrα-CreERT2;taumGFP* mice dosed with tamoxifen at P60 as previously described^10^. We used tissue clearing and light-sheet 3D imaging of the tau-mGFP signal, which in this model localises to adult-generated myelin. Strikingly, new myelin formation was restricted to the ME in the hypothalamus and absent from the paraventricular nucleus of the hypothalamus and the supraoptic nucleus, where magnocellular AVP cell bodies reside (**Fig. 5p, Suppl. Video 1**). Thus, myelin turnover on AVP axons is restricted to their terminal segments at the level of the ME.

### Adult oligodendrogenesis regulates ADAMTS4 expression and extracellular matrix composition in the MBH

The absence of glycaemic dysfunction in *Mbp^fl/fl^* mice indicates that intact myelination is not required for normal glucose homeostasis and that reduced myelination of ME axons in *Myrf^fl/fl^* mice does not underpin their glycaemic phenotype. Thus, we considered alternative mechanisms through which adult OL differentiation might regulate neuroendocrine and glucoregulatory functions. We previously found that production of new OLs in adulthood contributes to the remodelling of perineuronal nets (PNNs)^11^. PNNs are glycosaminoglycan (GAG)-rich extracellular matrix (ECM) structures that enmesh neurones at the ME-ARH border^21^ and have been implicated in the regulation of glucose homeostasis^22^. Key components of PNN scaffolds as well as enzymes regulating their post-translational modification are enriched in newly formed OLs^11^, which are abundant in the ME. Thus, we tested the hypothesis that impaired PNN organisation in the ME of *Myrf^fl/fl^*mice might produce their neuroendocrine and glycaemic phenotype.

We first assessed the role of adult oligodendrogenesis in the regulation of MBH PNN density. Visualisation of chondroitin sulphate (CS)-GAGs using *Wisteria floribunda* agglutinin (WFA) lectin demonstrated a significant reduction in the density of WFA^+^ PNNs in the ME and ARH of *Myrf^fl/fl^* mice (**Fig. 6a-b**), indicating that new OLs regulate PNN density here. To further evaluate how genetic block of new OL production might lead to these changes, we measured the relative gene expression of PNN structural components expressed by ME OL lineage cells^11^ including CSPGs, glycoproteins, link proteins and enzymes mediating PNN post-translational modifications. None of these transcripts were differentially expressed in *Myrf^fl/fl^* mice compared to littermate controls, except for *Adamts4* (*A disintegrin and metalloproteinase with thrombospondin motifs 4*) which was strikingly absent from the ME of *Myrf^fl/fl^* mice (**Fig. 6c-d**). Lack of *Adamts4* transcript in the ME of *Myrf^fl/fl^* mice was associated with a robust decrease in ME ADAMTS4 expression (**Fig. 6f**).

**Figure 6:**
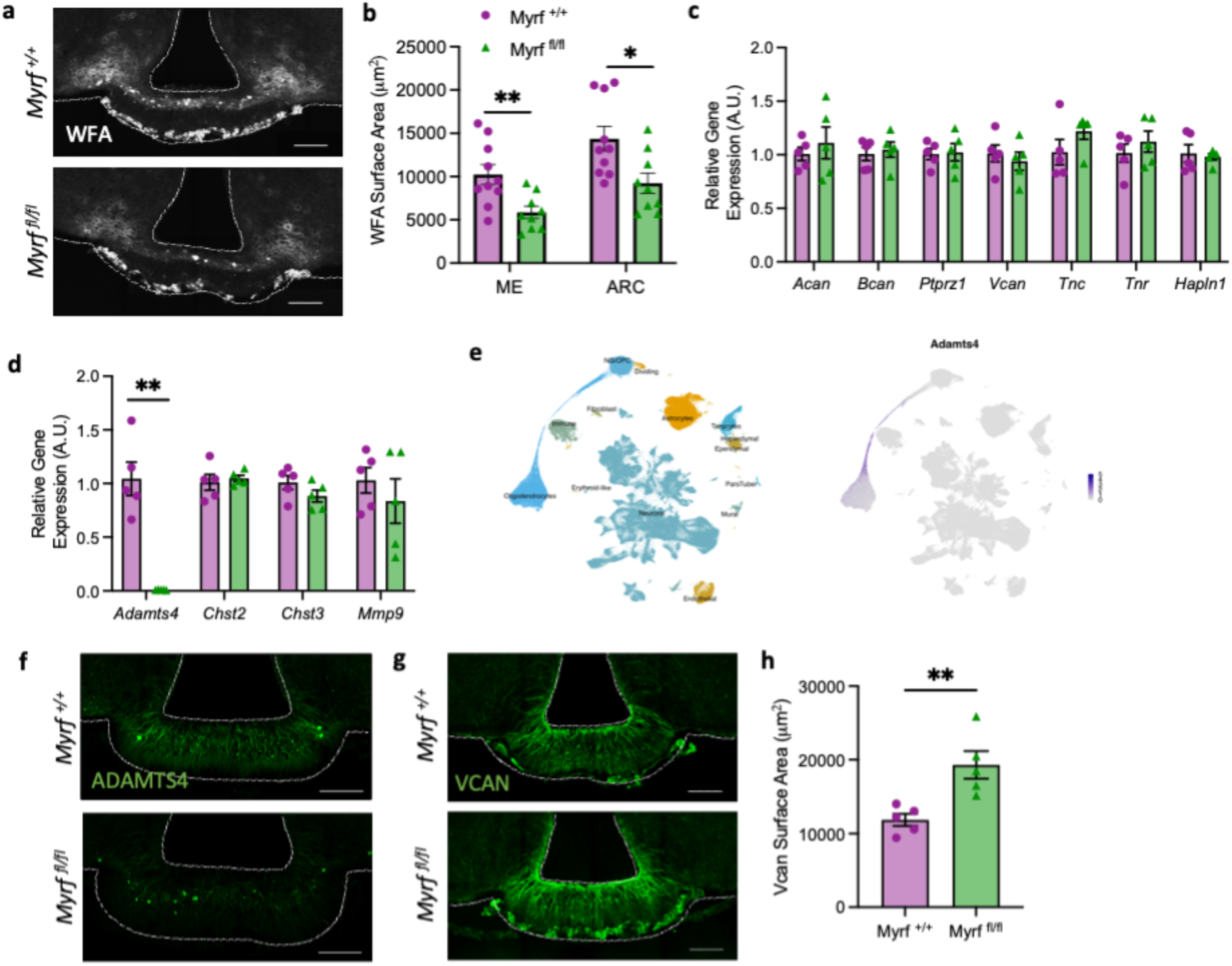
Oligodendrocyte progenitor cell differentiation regulates perineuronal net density and composition in the mediobasal hypothalamus. **(a)** Representative images of *Wisteria floribunda agglutinin* (WFA) lectin labelling in the median eminence (ME) and arcuate nucleus (ARC) of myelin regulatory factor (MYRF) conditional knockout mice (*Myrf ^­­^*) and controls (*Myrf ^­­^*) six week after tamoxifen administration at postnatal day 60 and (**b**) associated quantification. Gene expression analysis of transcripts relating to (**c**) perineuronal net (PNN) structural components and (**d**) PNN-modifying enzymes expressed by oligodendrocyte lineage cells in the mediobasal hypothalamus of *Myrf ^­­^*and *Myrf ^­­^*mice. (**e**) (left) Hypothalamic cell populations from HypoMap^­­^. (right) Log-normalized expression of *Adamts4* in the hypothalamic cell populations, demonstrating specific expression in OL populations. (**f**) Representative images of ADAMTS4 labelling in the ME-ARC of *Myrf ^­­^*and *Myrf ^­­^*mice. (**g)** Representative images of Versican (VCAN) labelling in the ME-ARC of *Myrf ^­­^*and *Myrf ^­­^*mice and (**h**) associated quantification. Data presented as mean ± S.E.M. and analysed by unpaired student’s - or Welch’s – t test, *p<0.05, **p<0.01, scale bars represent 100 μm, n=5-10/group.

ADAMTS4 is a metallopeptidase important for ECM and vascular remodelling through its role in CSPG degradation^23,24^. Transcriptomic data indicate that brain *Adamts4* is expressed specifically by OLs^25,26^ and enriched in OLs undergoing active myelination^27^, suggesting that *Myrf* knockout leads to ablation of the OL subset expressing *Adamts4*. To further understand the molecular identity of the OL subset expressing *Adamts4* in the ME, we analysed the expression of *Adamts4* in the HypoMap^19^, a publicly available transcriptomic reference map of the murine hypothalamus. This demonstrated that hypothalamic *Adamts4* expression is restricted to OL lineage cells (**Fig. 6e, Extended Data Fig. 5**), where it is enriched in newly formed myelinating OLs (**Fig. 6e, Extended Data Fig. 5**). Versican (VCAN) is a primary target of ADAMTS4^28^ and is highly expressed by ME OL lineage cells^11^. Consistent with decreased *Adamts4* expression, VCAN immunolabelling was significantly increased in *Myrf^fl/fl^* mice compared to controls (**Fig. 6g-h**), suggesting decreased degradation of VCAN in this model.

### ADAMTS4 regulates the MBH ECM, vascular permeability and glucose homeostasis

To further define the role of decreased ADAMTS4 expression in the changes observed in *Myrf^fl/fl^* mice, we examined hypothalamic tissues from *Adamts4* knockout (*Adamts4^-/-^*) mice. We confirmed absence of ADAMTS4 expression in the ME (**Fig. 7a**), which was associated with an upregulation of VCAN immunolabelling (**Fig. 7b-c**) and a decrease in WFA labelling in the ME and ARH (**Extended Data Fig. 6a-b**). ME MBP expression was unaffected by *Adamts4* knockout (**Extended Data Fig. 6c-d**) but was associated with increased BCAS1 expression (**Extended Data Fig. 6e-f**), indicating that *Adamts4* deletion does not impair new OL generation and myelination in the ME. However, *Adamts4* knockout significantly increased the number of MECA32+/Laminin+ fenestrated capillary loops in the ventral ME (**Fig. 7d-e**) indicating a role for ADAMTS4 in ME vascular remodelling. Consequently, we used albumin immunostaining to assess permeability of the blood-hypothalamus barrier in this model^29^. Consistent with increased density of fenestrated vessels, we found that albumin diffusion within the ME and ARH was significantly increased in *Adamts4^-/-^* mice (**Fig. 7f-g**). Thus, *Adamts4* deletion is sufficient to increase vessel fenestration and vascular permeability of the blood-hypothalamus barrier. Consistent with the glycaemic phenotype of *Myrf^fl/fl^* mice, blood glucose and plasma corticosterone levels were significantly increased in *Adamts4^-/-^* mice compared to controls (**Fig. 7h-i**), supporting a role for ADAMTS4 in the regulation of glucose homeostasis.

**Figure 7:**
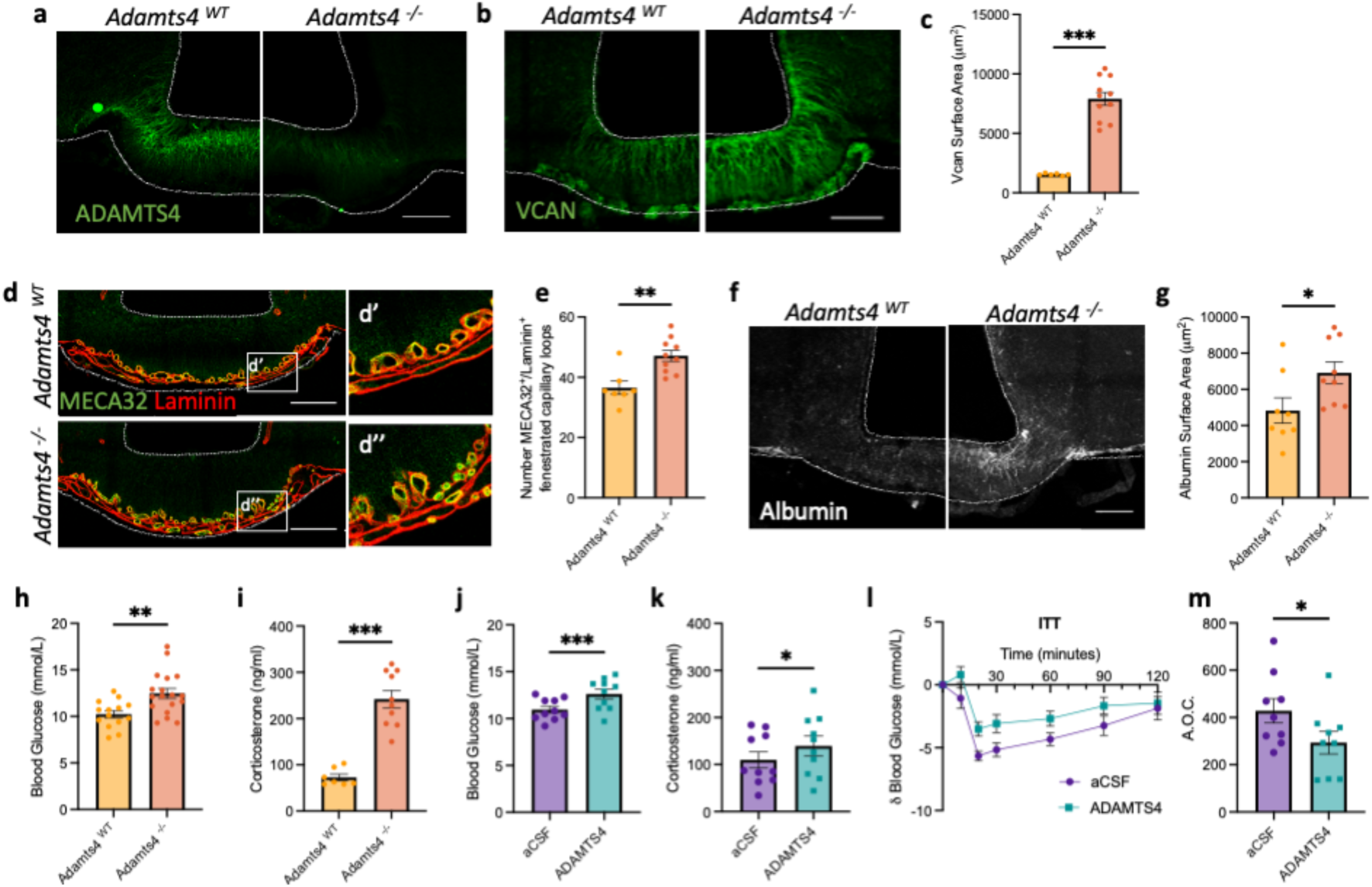
ADAMTS4 regulates the extracellular matrix and vascular permeability in the mediobasal hypothalamus. (**a**) A disintegrin and metallopeptidase with thrombospondin motifs 4 (ADAMTS4) expression in the mediobasal hypothalamus (MBH) of ADAMTS4 knockout (*Adamts4^­­^*) mice compared to controls (*Adamts4^­­^*). Representative images of (**b**) versican (VCAN), (**d**) MECA32+/laminin+ fenestrated capillary loops and (**f**) albumin immunolabelling in *Adamts4^­­^*and *Adamts4^­­^*mice and associated quantifications (**c**, **e**, **g**). (**h**) Blood glucose and (**i**) plasma corticosterone levels in *Adamts4^­­^*and *Adamts4^­­^*mice at ZT2. (**j**) Blood glucose, (**k**) plasma corticosterone and (**l-m**) insulin sensitivity in C57BL/6J mice one hour after intra-hypothalamic aCSF or ADAMTS4 administration. Data presented as mean ± S.E.M. and analysed by unpaired student’s t-test or Mann-Whitney test, *p<0.05, **p<0.01, ***p<0.001, ****p<0.0001, n=8-18/group. Scale bars represent 100 μm.

Next, we performed bilateral injections of artificial cerebrospinal fluid (aCSF) or mouse recombinant ADAMTS4 directly into the MBH of adult C57BL/6J mice via a pre-implanted cannula. Consistent with a role for ADAMTS4 in VCAN degradation, ADAMTS4 injection rapidly decreased VCAN immunolabelling in the MBH (**Extended Data Fig. 6g-h**). However, the density of fenestrated capillary loops in the ME of ADAMTS4 injected animals was unchanged after one hour (**Extended Data Fig. 6i-j**). Moreover, against our prediction, MBH ADAMTS4 administration also increased blood glucose levels (**Fig. 7j**) and plasma corticosterone (**Fig. 7k**) within 1 hour following injection, and significantly decreased insulin sensitivity (**Fig. 7l-m**), suggesting a bimodal role of ADAMTS4 in the regulation of blood glucose and corticosterone levels.

### ME ADAMTS4 expression is downregulated during hypoglycaemia

Our results indicate that ME adult oligodendrogenesis rapidly responds to changes in blood glucose levels and regulates ADAMTS4 expression and vascular permeability. To determine whether acute and chronic changes in blood glucose levels are sufficient to change ADAMTS4 expression by ME OLs, we examined ME ADAMTS4 expression following acute insulin-induced hypoglycaemia or in diabetic mice. Insulin-induced hypoglycaemia significantly decreased ADAMTS4 expression (**Fig. 8a-b**), reduced the density of WFA+ ECM in the ME (**Extended Data Fig. 7a-b**) and increased ME VCAN expression (**Extended Data Fig. 7c-d**), suggesting that ADAMTS4 is regulated by acute changes in peripheral glycaemia. We then used a model of diabetes, achieved through the combination of diet-induced-obesity (DIO) and streptozotocin (STZ) administration in C57BL/6J mice to induce insulin resistance and pancreatic β cell dysfunction. Compared to vehicle treated animals (DIO-Veh), treatment with STZ (DIO-STZ) increased *ad libitum* fed (**Extended Data Fig. 7e**) and fasting (**Fig. 8c**) blood glucose and produced glucose intolerance (**Extended Data Fig. 7f-g**). Chronic hyperglycaemia in this model was associated with increased ADAMTS4 expression in the ME (**Fig. 8e-f**). In addition, we found a strong correlation between blood glucose levels and ME ADAMTS4 expression (**Fig. 8g**). Thus, changes in blood glucose levels regulate the expression of ADAMTS4 by oligodendrocytes.

**Figure 8:**
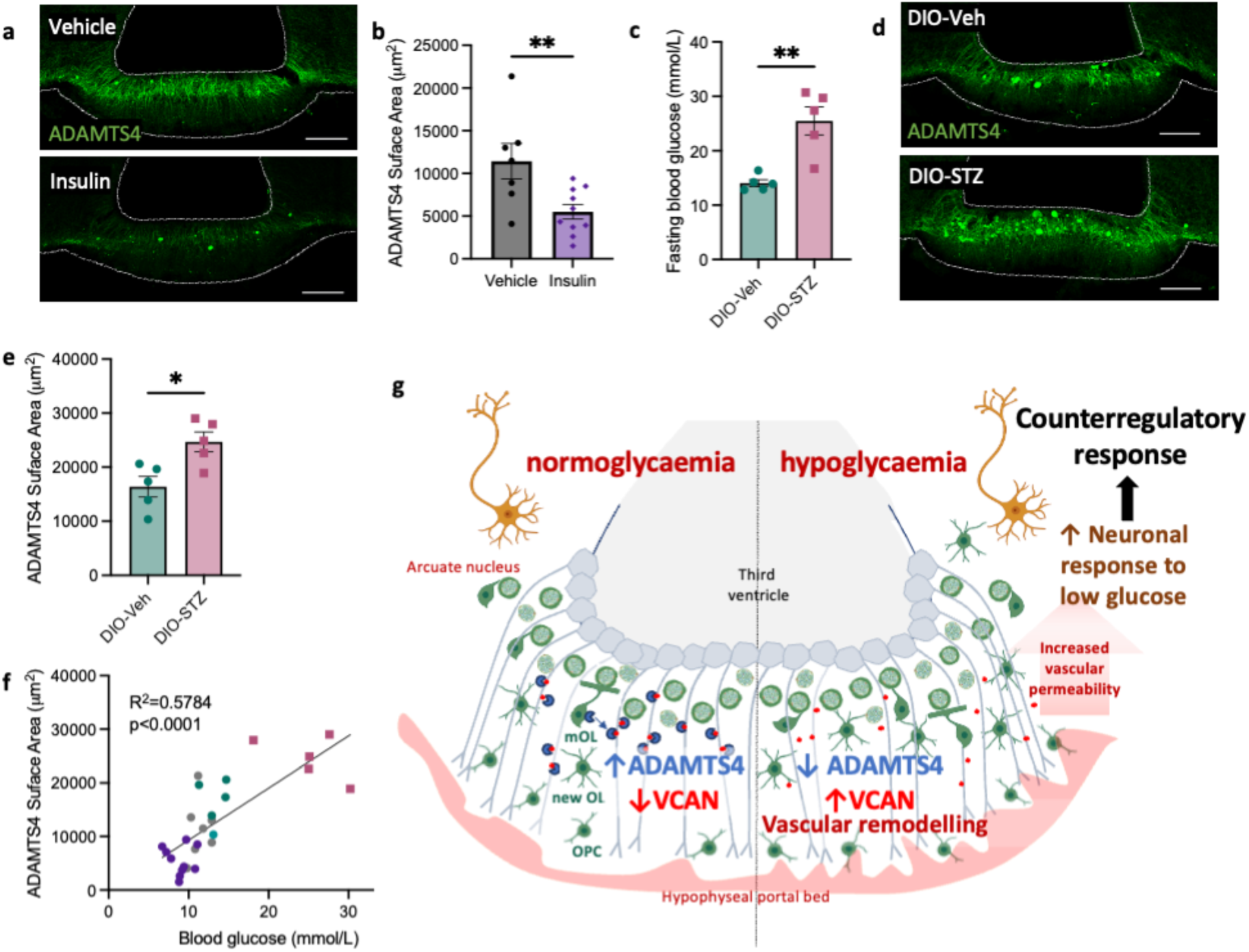
Median eminence ADAMTS4 expression is regulated by glycaemia. Representative images of A disintegrin and metalloproteinase with thrombospondin motifs 4 (ADAMTS4), immunolabelling in the mediobasal hypothalamus (MBH) (**a**) one hour after intraperitoneal administration of vehicle (saline; 10 ml/kg) or insulin (0.75 U/kg) and (**b**) associated quantification. (**c**) Blood glucose levels in mice fasted for 6 hours in diet-induced-obese (DIO) mice four weeks after administration of vehicle (DIO-Veh) or streptozotocin (DIO-STZ). (**d**) ADAMTS4 immunolabelling in the MBH of DIO-Veh and DIO-STZ mice and (**e**) associated quantification. (**f**) Correlation between ME ADAMTS4 immunolabelling and blood glucose across multiple models, analysed by linear regression. (**g**) Working model by which ME OL lineage cells regulate glucose homeostasis. All data presented as mean ± S.E.M., scale bars represent 100 μm. Data analysed using an unpaired student’s- or Welch’s-t test, *p<0.05, **p<0.01, n=5-10/group.

## Discussion

Here, we identify a role for adult oligodendrogenesis in the central response to hypoglycaemia and the activation of the counterregulatory response that restores glucose homeostasis. We show that changes in blood glucose levels regulate the density of newly formed oligodendrocytes, which in turn regulate blood-hypothalamus barrier permeability, neuronal glucose sensing in the ARH, and neuroendocrine axes regulating blood glucose levels. We implicate oligodendrocytic expression of ADAMTS4 in these responses and show that expression of ADAMTS4 in the ME bidirectionally responds to physiological changes in blood glucose levels (**Fig. 8g**).

Our data support accumulating evidence that ME OPCs rapidly differentiate into new OLs in response to nutritional signals^10,11^. Until now, the exact nutritional signals that modulate ME OPC proliferation and differentiation in adulthood have remained unknown. Our results indicate that hypoglycaemia increases new OL production in the ME independently of circulating insulin levels and in the absence of changes in OPC proliferation or number. Instead, BCAS1 expression and the density of OL lineage cells in the ME are increased, suggesting that hypoglycaemia promotes survival of newly formed OLs. In contrast during 2DG-induced hyperglycaemia, ME OPC proliferation and density are increased but this increase is offset by a decrease in the density of newly formed OLs. This observation contrasts our previous report that refeeding promotes ME OPC differentiation^11^. In that context, postprandial hyperglycaemia is accompanied by numerous other changes in circulating nutrients and hormones, which might collectively change the outcome on OPC differentiation and survival of newly formed OLs. In addition, 2DG-induced hyperglycaemia is sustained and more severe than what is observed physiologically after a meal. Thus, the observed decrease in the density of newly formed OLs might reflect a consequence of glucotoxicity.

Our work identifies new OL production as a regulatory component of the ME vascular and diffusion barriers. Genetic block of new OL production phenocopies some of the vascular and structural alterations observed in the ME during fasting (increased vascular permeability and decreased tanycytic paracellular permeability) that were previously shown to be mediated by tanycytes^6^. This suggests that multi-cellular mechanisms are recruited to mediate the remodelling of the ME vasculature during changes in energy or nutrient availability. Blockade of new OL generation also impairs the ME-ARH barrier, increasing the access of blood-borne signals to the ARH with functional consequences on local neurocircuit activity. This contrasts with previous work indicating that tanycytes are not required for the maintenance of the ME-ARH barrier in a model of tanycyte ablation^30^. Consistently, increased ME-ARH permeability in *Myrf* conditional knockout (cKO) mice occurs in the absence of changes in tanycytic density and morphology. Thus, OL plasticity is required for the maintenance of the diffusion barrier between the ME and ARH, a function that does not require an intact tanycytic population.

In the context of 2DG-induced neuroglycopenia, we observe that adult oligodendrogenesis regulates MBH glucose sensing. *Myrf* cKO mice exhibit an accelerated and augmented hyperglycaemic response to neuroglycopenia, suggesting that changes in the ME-ARH barrier regulate both the speed and amplitude at which MBH circuits respond to hypoglycaemia and recruit the HPA axis. Thus, these findings support a role for ME-ARH glucose sensing mechanisms in the hypothalamic response to hypoglycaemia, which have, so far, mainly been attributed to glucose sensors located in the VMN. How these distinct sensors collaborate to produce the counter-regulatory response to hypoglycaemia remains to be evaluated but might be recruited sequentially during exposure to hypoglycaemia. Indeed, ME sensors are neuroanatomically well positioned to rapidly detect changes in circulating glucose levels and engage counterregulatory mechanisms that restore glucose homeostasis^31^. Well-established glucose sensors in the ME-ARH region include AgRP neurones, although their role in the counterregulatory response to hypoglycaemia needs further evaluation. AgRP neurons modulate the activity of the HPA axis via both projections to PVH CRH cell bodies^32,33^ and direct projections to CRH terminals in the ME, providing a local mechanism for rapid activation of CRH release^34^. In contrast, whether VMN glucose sensors can rapidly respond to decreased circulating glucose levels is unclear^31^. These parenchymal sensors might be engaged at a later stage following remodelling of the ME-ARH barrier, which would expose them to decreased interstitial glucose concentrations. While we and others^6^ demonstrate that structural reorganisation of the ME is a component of the response to hypoglycaemia, how quickly these structural changes occur and whether they allow differential access to the VMN in a short timeframe remains to be examined.

Strikingly, our results indicate that adult oligodendrogenesis contributes to the neuroendocrine response to hypoglycaemia independently of myelination. Conditional *Mbp* deletion in adult OPCs, which maintains OPC differentiation, but blocks the formation of new compact myelin, does not impair glucose tolerance or the counterregulatory response to hypoglycaemia. This is consistent with an acute role for adult oligodendrogenesis in glycaemic control since acute changes in blood glucose levels do not change ME myelination. Magnocellular axons which pass through the ME and are myelinated have been implicated in the regulation of pancreatic glucagon release^20^. However, plasma glucagon levels at baseline or following a 2DG challenge did not differ between *Myrf* cKO mice and controls. In addition, decreased myelination of magnocellular axons in the ME of *Myrf* cKO mice fails to impair osmoregulation during dehydration, an important functional output of this axis. These results support the conclusion that the output of hypothalamic magnocellular neurons is not altered in *Myrf* cKO mice. This is consistent with our observation using whole brain 3D visualisation of newly formed myelin that myelin plasticity is strictly localised to the ME (at least in the timeframe of these studies) and not observed along magnocellular axons outside the ME. Thus, we infer that local ME demyelination does not impinge on the glucoregulatory actions of AVP neurones under our experimental context.

Our study instead implicates that pre-myelinating OLs contribute to the neuroendocrine response to hypoglycaemia through ADAMTS4, a metallopeptidase that is expressed exclusively within the OL lineage, enriched in newly differentiating OLs^11,19,26^ and robustly downregulated in the ME of *Myrf* cKO mice. In line with reports of insulin-induced inhibition of ADAMTS4 expression in fibroblast-like synoviocytes^35^ and genome wide association studies demonstrating an association between human ADAMTS4 variants and fasting insulin levels^36^, we find that MBH ADAMTS4 availability is physiologically regulated by acute changes in glycaemia and becomes dysregulated in diabetic mice. Furthermore, we demonstrate that ADAMTS4 regulates the HPA axis, glucose homeostasis and ME vascular permeability. Notably, *Adamts4* depletion in *Myrf* cKO mice and administration of exogenous ADAMTS4 into the MBH of wild-type mice both result in insulin resistance and hypercorticosteronaemia. However, the mechanism(s) by which both increased and diminished ADAMTS4 availability influence glucose homeostasis and the HPA axis in the same direction are unclear. Of note, while endogenous ADAMTS4 expression is restricted to the ME and absent from the ARH, we likely produced significant PNN remodelling in the ARH following pharmacological administration of ADAMTS4 into the MBH. ADAMTS4 has a potent aggrecanase activity^37^ and aggrecan is present in ARH^21^. Thus, the response observed following exogenous ADAMTS4 injection might be confounded by the loss of aggrecan in the ARH.

*Adamts4* knockout phenocopies the alterations observed in the ME of *Myrf* cKO mice, indicating a direct role for ADAMTS4 in the regulation of blood-hypothalamus barrier permeability. Previous work implicates ADAMTS4 in the regulation of angiogenesis *in vitro* through the sequestration of vascular endothelial growth factor (VEGF)^38^, a factor upregulated by ME tanycytes in response to hypoglycaemia^6^. One mechanism by which oligodendrocytic ADAMTS4 might regulate ME vascular permeability is via its interaction with VEGF. Alternatively, ADAMTS4 might regulate vascular permeability via effects on VCAN density, of which the V2 isoform (specifically measured in our study) displays angiogenic properties *in vitro*^39^. Our data demonstrate that physiological and experimental manipulations of ME ADAMTS4 expression go hand in hand with changes in VCAN, with ADAMTS4 expression inversely associated with ME VCAN amounts. In insulin-treated animals, for example, ADAMTS4 is rapidly downregulated and is associated with increased ME VCAN. Since loss of *Adamts4* in *Myrf* cKO mice is not associated with altered *Vcan* transcript expression we infer that ADAMTS4 regulation of ME VCAN amounts occurs at the post-translational level i.e. via altered degradation, providing an additional mechanism by which adult oligodendrogenesis might rapidly regulate ME vascular permeability.

Counter-intuitively we observe that WFA immunolabelling in the MBH decreases in *Adamts4* knockout mice despite an increase in VCAN amounts. Since the WFA lectin only binds to neutrally charged CS-GAGs^40^, the decrease in WFA+ PNNs in this model may reflect changes in the charge of local CS-GAGs, for example, through changes in glycosylation or sulphation. Future studies should therefore address how OL differentiation modulates post-translational processing of CS-GAGs, as this may uncover additional mechanisms by which OLs regulate local barrier properties.

Finally, our finding that ADAMTS4 expression becomes dysregulated in diabetic rodents is noteworthy and warrants further investigation. Therefore, a relevant question for future studies is to define whether OL-derived ADAMTS4 plays a causal role in metabolic dysfunction associated with obesity and diabetes via effects on blood-brain-barrier integrity^41^ and/or ME-ARH PNN composition and density^21^.

## Methods

### Animals

All animal experiments were performed in accordance with the UK Home Office regulations under the Animals (Scientific Procedures) Act (1986) and with the approval of the University of Cambridge Animal Welfare and Ethics Review Board. Animals were group-housed in a specific pathogen free facility and maintained on a standard 12-hour light/dark cycle (lights on 7:00-19:00) at 22°C with *ad libitum* access to water and standard laboratory chow (SAFE R105, SAFE Complete Care Competence, Rosenberg, Germany) unless otherwise stated. All experiments were performed on male mice starting from postnatal day 60 (P60). C57BL/6J mice were obtained from Charles Rivers Laboratories (Saffron Walden, UK). *Myrf^fl/fl^* mice^42^ and *Mbp**^fl/fl^***mice (S.G.N., H.L. and W.D.R. unpublished) were provided by Professor William Richardson at University College London. Note that this *Mbp^fl/fl^* line is structurally and phenotypically like that described by Meschkat et al.^43^ but was conceived and made independently. Tissues from *Adamts4^-/-^* mice were obtained from Professor Satoshi Hirohata at Okayama University.

### Tamoxifen administration and preparation

Tamoxifen (Sigma) was prepared in corn oil (Sigma) by sonication at 37°C at 30 mg/ml prior to administration by oral gavage at 300 mg/kg of body weight on 4 consecutive days.

### Glucose challenges in C57BL/6J mice

C57BL/6J mice were randomly allocated to groups and received four intraperitoneal (i.p.) injections of BrdU (Sigma; 50 mg/kg - prepared in sterile saline at 5 mg/ml) within a 24-hour period starting at ZT5 and ending at ZT4 on the following day. Mice were fasted for 4 hours prior to the fourth injection, when mice received BrdU alone (vehicle) or in combination with glucose (2 g/kg), insulin (0.75 U/kg) or 2DG (250 mg/kg). One hour later a blood glucose reading was obtained from the tail vein at ZT5 immediately prior to sacrifice using an AlphaTrak 2 handheld glucometer (Precision Xtra; MediSense).

### Streptozotocin-induced diabetes in diet-induced-obese mice

C57BL/6J mice were fed a high fat diet with 60% kilocalories obtained from fat (60% HFD; Research Diets D1492i) from 8 weeks of age for 8 weeks. After 8 weeks, animals were randomly allocated to groups and a baseline blood glucose reading was obtained from the tail vein using an AlphaTrak 2 handheld glucometer prior to ip administration of either vehicle (saline; 10 ml/kg) or streptozotocin (STZ, Sigma; 100 mg/kg). Metabolic phenotyping tests were performed 4 weeks after vehicle or STZ administration.

### Hyperinsulinaemic clamp paradigm

In brief, C57BL/6J mice underwent surgery under isoflurane anaesthesia using a low-flow anaesthesia delivery system (SomnoSuite® - Kent Scientific Corporation) to implant vascular catheters into the left carotid artery and right jugular vein. Vascular lines were exteriorised behind the nape of the neck using a vascular access button (Instech Laboratories) allowing post-operative group-housing.

On study days, animals were fasted from ZT1 and vascular lines flushed and connected via a tether and low-torque dual channel swivel to allow handling-free intravenous infusion and arterial blood sampling. After acclimatization, starting around midday (ZT5), a non-primed continuous intravenous infusion (10 mU/kg/min) of rapid-acting insulin (NovoRapid, Novo Nordisk diluted in Bovine Serum Albumin, Sigma-Aldrich) was delivered together with a simultaneous infusion of dextrose (20% (w/v) or 50% glucose (w/v)-Fresenius Kabi and Hameln pharma respectively). Arterial samples to measure whole blood glucose concentrations were taken every 5-15 minutes and analysed using a handheld blood glucose meter (Accu-Chek® Aviva, Roche) to minimize blood volume sampling. Additional samples were taken periodically for comparison with plasma glucose concentrations determined biochemically using an Analox Analyser (GM9 Glucose Analyser, Analox Instruments). Dextrose infusion rates were adjusted to achieve target ranges of ≤ 3.9, 6-8 and ≥ 15.0 mmol/L for hypoglycaemia, euglycaemia and hyperglycaemia respectively. Clamps were 75 minutes in length, to allow roughly 30 minutes to reach the target glycaemic level, and then 45 minutes in a steady state. If necessary, clamps were extended until blood glucose had been stable within the target range for at least 40 minutes. Plasma samples for insulin analysis were collected at baseline and at the end of the clamp. Upon completion of the clamp, mice immediately underwent perfusion fixation.

### Metabolic tolerance tests

Mice were moved to a procedure room at ZT1, singly housed and fasted in clean cages with home cage enrichment. A baseline blood glucose reading was taken from the tail vein using a handheld glucometer at ZT5. Mice were administered an oral gavage of glucose (2 g/kg; Sigma) for an OGTT, an ip injection of glucose (1 g/kg; Sigma) for an ipGTT or an ip injection of insulin (0.75 U/kg; Actrapid, Novo Nordisk) for an ITT at ZT6 and blood glucose measured 10, 20, 30, 60, 90 and 120 minutes later. For OGTTs, additional blood samples were obtained at baseline and 20 minutes after glucose administration for measurement of plasma insulin. At the end of the test, mice were returned to home cages and refed.

### Dehydration paradigm

Water was removed from the animals’ home-cage immediately prior to the dark onset (ZT12). Plasma samples were obtained at baseline and after 12 hours water deprivation (ZT0) for measurement of plasma osmolarity using an Osmomat 3000 (Gonotec). Following plasma sampling, water was returned to the home cage.

### 2DG studies

Glucoprivic feeding and hyperglycaemia were assessed via crossover studies. For all experiments, mice were moved to a procedure room, singly housed, and fasted at ZT1 prior to i.p. injection of either saline (Vetivex) or 2DG (250 mg/kg; Sigma) at 10 ml/kg of body weight at ZT5. For food intake studies, animals were provided with standard chow diet 30 minutes after saline or 2DG administration and food intake measured at 30, 60, 90, 120 and 180 minutes as loss of food weight in grams. For blood glucose measurements, blood glucose readings were obtained from the tail vein using a handheld glucometer 0, 15, 30, 60, 90 and 120 minutes after saline or 2DG injection. For measurement of counter-regulatory hormones, plasma samples were collected 30 minutes after saline or 2DG administration. For assessment of 2DG-induced cFos expression, mice were fasted from ZT1, administered saline or 2DG at ZT5 as described above and underwent perfusion-fixation 90 minutes later.

### Plasma analyses

Whole blood samples were obtained from the tail vein into heparinised capillary tubes and kept on ice until centrifugation with a Haematospin 1300 (Hawksley) to obtain plasma. Glucagon was measured using an enzyme-linked immunosorbent assay (Mercodia AB; product code 10-1271-01). Insulin was measured using the Mouse/Rat Insulin Kit on the MesoScale Discovery immunoassay platform (MesoScale Delivery; product code K152BZC-3). ACTH was measured by sandwich immunoassay using the MilliPlex MAP Mouse Pituitary Magnetic Bead Panel Kit (Merck Millipore; product code MPTMAG-49K). Corticosterone was measured using the Corticosterone EIA Kit (ImmunoDiagnosticSystems; product code AC14F1).

### ADAMTS4 injection paradigm

Surgical procedures were carried out in single-housed C57BL/6J mice under isoflurane anaesthesia that received Metacam prior to surgery. Steel guide cannula (Plastics One) were stereotactically implanted 1 mm above the ME-ARH region (A/P: - 1.3 mm, D/V: - 5.0 mm, lateral: + 0.2 mm relative to Bregma) and secured using Loctite glue and dental cement (Fujicem). All mice were acclimatised to handling and the brain injection procedure during the one-week post-operative period. Following recovery from surgery, injections were performed in a crossover manner so that all mice received aCSF on one week and one week later received ADAMTS4. Where injections were repeated, mice were given at least a 4-week wash out period between ADAMTS4 and aCSF injections. On the day of injection, mice were moved to a procedure room at ZT1 and fasted in clean cages with home cage enrichment. Bilateral injections of aCSF (Tocris) or 4 ng recombinant human ADAMTS4 (R&D Systems 4307-AD; 2 ng/side in 100 nl) were performed at ZT5 using bevelled stainless steel 33-gauge injectors, extending 1 mm beyond the tip of the guide. Plasma samples were collected from the tail vein one hour after aCSF or ADAMTS4 injection at ZT6, after which mice were refed and returned to the holding room for daily body weight and food intake measurements in the week after injection. Insulin tolerance tests (ITTs) were started one hour after aCSF/ADAMTS4 administration at ZT6 as described above.

### Perfusion fixation

Mice were administered 50 ul pentobarbital (Dolethal, 200 mg/ml) by intraperitoneal injection to achieve deep anaesthesia. Mice were then trans-cardially perfused with 50 ml heparinised phosphate buffered saline (PBS) followed by 4% (w/v) paraformaldehyde (PFA; Fisher Scientific, Waltham, Massachusetts) in PBS (pH 7.4) or 4% PFA with 2.5% (v/v) glutaraldehyde (Sigma) in 0.1 M phosphate buffer (pH 7.4) at a flow rate of 5 ml/min. For BBB permeability assessments using Evans blue, mice were perfused with heparinised PBS followed by 4% PFA containing 1% Evans blue (w/v) at a flow rate of 7 ml/min. Following perfusion, brains were dissected and immersed in the appropriate fixative. PFA-fixed brains for immunofluorescent staining or cFos immunohistochemistry were transferred to 30% sucrose (w/v; Sigma) in PBS after 24-48 hours in fixative.

## Tissue Processing

### Sectioning of PFA-fixed tissues

Following cryoprotection in 30% sucrose (w/v; Sigma) in PBS, PFA-fixed brains were embedded in Optimal Cutting Temperature (OCT; CellPath) and sectioned at 20 – 30 μm in the coronal plane from Bregma -1.58 to -2.30 mm^44^ using a Leica SM2010R Sliding Microtome or Leica CM1950 Cryostat (Leica Biosystems). Except for sections obtained from mice with bilateral cannula, all sections were collected into cryoprotectant solution and processed as free-floating sections.

### Immunofluorescent staining

Sections were washed in PBS prior to heat mediated antigen retrieval in 20 mM sodium citrate buffer (pH 6.5; Fisher Scientific) for 20 minutes at 80°C on a heat block. Where sections were immunolabelled for BrdU, sections were subjected to an additional antigen retrieval step in 2 N hydrochloric acid (Fisher Scientific) for 30 minutes in a water bath pre-heated to 37°C followed by incubation with 0.1M sodium borate buffer (pH 8.5; Fisher Scientific) for 10 minutes at room temperature. Sections were then washed in PBS prior to blocking with 3-10% (v/v) normal donkey – or goat-serum (NDS/NGS; Abcam) in PBS with 0.3% (v/v) Triton X-100 (0.3% PBST) for one hour at room temperature with agitation. Primary antibodies were prepared in blocking solution and incubated with sections for 1-2 nights at 4°C with agitation. Sections were washed in 0.1% PBST prior to incubation with secondary antibodies diluted at 1:500 in 0.3% PBST for 2 hours at room temperature. Antibodies targeting different antigen were multiplexed where possible using a sequential staining approach. Following staining, sections were mounted to slides under coverslips with VectaShield HardSet Antifade Mounting Medium with DAPI (Vector Laboratories), left to cure overnight at room temperature and stored at 4°C until imaging.

Primary antibodies were goat anti-Sox10 (R&D Systems AF2864, 1:50), rabbit anti-PDGFRa (Cell Signalling Technologies 3164, 1:500), rat anti-BrdU (Abcam ab6326, 1:200), guinea pig anti-Bcas1 (Synaptic Systems 445 004, 1:500), rabbit anti-Bcas1 (Synaptic Systems 445 003, 1:1000), guinea pig anti-opalin (Synaptic Systems 457 005, 1:500), rat anti-MBP (Abcam ab7349, 1:500), chicken anti-GFP (Abcam ab13970, 1:1000), Wisteria floribunda agglutinin lectin (Vector Laboratories B-135, 1:500), rabbit anti-versican (Sigma-Aldrich AB1032, 1:500), rabbit anti-ADAMTS4 (Invitrogen PA1-1749A, 1:500), chicken anti-vimentin (Millipore AB5733, 1:1000), rabbit anti-Zo-1 (Invitrogen 61-7300, 1:250), rabbit anti-laminin (Sigma-Aldrich L9393, 1:500) and rat anti-MECA32 (NovusBio NB100-77668, 1:50).

Secondary antibodies were biotinylated goat anti-rabbit (Vector Laboratories BA-1000-15), goat anti-guinea pig 647 (ThermoFisher Scientific A21450), goat anti-rabbit 555 (ThermoFisher Scientific A21428), goat anti-rat 488 (ThermoFisher A11006), donkey anti-chicken 488 (Jackson ImmunoResearch 703-545-155), donkey anti-goat 647 (ThermoFisher Scientific A21447), donkey anti-rabbit 488 (ThermoFisher Scientific A21206), donkey anti-rabbit 555 (ThermoFisher Scientific A31572), donkey anti-rabbit 594 (ThermoFisher Scientific A21207), donkey anti-rat 488 (ThermoFisher Scientific A21208), Streptavidin 594 (ThermoFisher Scientific S11227) and Streptavidin 647 (ThermoFisher Scientific S21374).

### Confocal microscopy and analysis

Researchers were blinded to the experimental condition during imaging and analysis. Sections were imaged as z-stacks on a Leica SP8 confocal microscope at intervals of 2.0-3.3 μm using a 40x oil objective with tile scanning to obtain information from the entire depth and area of the region of interest (ROI). During image acquisition, microscope settings and z-stack size identical within each experiment. Images were analysed using Fiji (ImageJ) software. For analysis, z-stacks were projected into a single image and the freehand tool was used to draw, define, and calculate the ROI area. Cell counts were performed using the Fiji manual cell counter tool. Where area was measured, the same threshold was applied to all images prior to measurement of the area or area fraction. ROI borders were determined using the Paxinos and Franklin Mouse Brain Atlas^44^.

### cFos Immunohistochemistry

Sections were washed in PBS prior to incubation with 0.5% hydrogen peroxide (H_2_O_2_) in MilliQ water (MQ-H_2_O) for 15 minutes at room temperature. Following washing with PBS, sections were blocked in 5% NGS in 0.3% PBST for 1 hour at room temperature and incubated with rabbit anti-cFos (Synaptic Systems 226 003) diluted 1:1000 in blocking solution for 2 nights at 4°C with agitation. Sections were washed in 0.1% PBST prior to incubation with biotinylated goat anti-rabbit secondary antibody (Vector Laboratories BA-1000-15) diluted 1:500 in 0.3% PBST for 1 hour at room temperature. ABC Reagent (Vector Laboratories PK-6100) was prepared as per the manufacturer’s instructions and, following washing with 0.1% PBST, was incubated with sections for 1 hour at room temperature. Sections were washed in 0.1% PBST prior to incubation with 3’3’-Diaminobenzidine (DAB; Sigma D8001) solution (15 ml PBS, 500 μl 5 mg/ml DAB in MQ-H_2_O, 50 ul 1% H_2_O_2_) for 5-10 minutes at room temperature with agitation. Sections were washed in PBS, mounted to slides and dehydrated using increasing concentrations of ethanol (Sigma) prior to mounting under coverslips with Pertex mounting media (Fisher Scientific). Images were acquired using an Axioscan A1 Slide Scanner (Zeiss) using a 20x air objective. Images were analysed in a blinded manner by manual cell counting within ROIs in Zen Software (Zeiss).

### RNAscope single molecule fluorescent in situ hybridisation

Perfusion-fixed brains were embedded in paraffin blocks and sectioned in the coronal plane at 5 μm on a Leica RM2235 microtome. Sections were dewaxed with xylene and 100% ethanol and washed prior to processing. Single molecule fluorescent *in situ* hybridisation was performed as previously described^11^ using RNAscope technology and a Leica Bond Rx instrument. Mouse *Bmp4* was labelled with a RNAscope 2.5 LS Probe (Mm-Bmp4-C2; ACD, 401308 C2) and detected with Opal 570 (Akoya Biosciences, OPI-001006). Sections were imaged using a spinning disk Operetta SLS (PerkinElmer) in confocal mode with a sCMOS camera and 40x automated water-dispensing objective. All slides were imaged with identical gain and laser power settings and analysed in Harmony software (PerkinElmer).

### Scanning electron microscopy (SEM)

Brains were post-fixed in 4% PFA with 2.5% (v/v) glutaraldehyde (Sigma) in 0.1 M phosphate buffer (pH 7.4) overnight before moving to 0.1M phosphate buffer. Brains were sectioned in the coronal plane on a Leica VT1000s vibratome to obtain 200 μm thick sections. Regions of interest were dissected out, washed 3 times in distilled water and incubated in 2% osmium tetroxide (Agar Scientific) with 1.5% potassium ferricyanide (Agar Scientific) for 2 hours at 4°C. Samples were then washed 3 times in distilled water and incubated with 2% uranyl acetate (Agar Scientific) overnight at 4°C. The next morning, samples were washed 3 times in distilled water prior to dehydration through a series of washes in ethanol as follows: 25% for 15 minutes, 50% for 15 minutes, 75% for 15 minutes, 90% for 15 minutes, 100% for 15 minutes repeated twice. Samples were then embedded in Epoxy 812 resin (Agar Scientific) as follows: 12 hours in a mixture of resin and 100% ethanol at a ratio of 1:3 respectively, 12 hours in a mixture of resin and 100% ethanol at a ratio of 1:1 respectively, 12 hours in a mixture of resin and 100% ethanol at a ratio of 3:1 respectively, 12 hours in 100% resin repeated once. Samples were then cured at 70°C for 48 hours prior to sectioning to 100 nm on a Leica UC7 Ultramicrotome using a diamond knife (Diatome – Ultra 45). Once cut, sections were floated and collected on glow discharged silicon wafer chips to be imaged. Imaging was performed using a TESCAN Clara Scanning Electron Microscope at a magnification of 10,000X using a 2.5 kV electron beam and 300 picoamps of current.

G-ratio was quantified for at least 50 transverse axons per animal using Fiji (ImageJ) software by measuring the inner diameter of an axon and dividing by the outer diameter of the axon (axon + myelin sheath). Axons were excluded from analysis if they exhibited an awkward shape such that the diameter could not be defined. For axons demonstrating an elongated shape, the g-ratio was quantified from measurements taken perpendicular to the axis producing the longest diameter^45^. Quantification was only performed for axons displaying compact myelin sheaths. Axons were quantified in randomly selected fields of view (at least 5 per brain) from different regions within the ME to ensure inclusion of axons from the entire region of interest, including the ventral, dorsal, medial and lateral portions of the ME.

### Whole brain clearing

All brains were washed in phosphate-buffered saline (PBS, pH 7.4) for 3 × 30 min at room temperature. Whole-mount staining for GFP was performed according to the previously described iDISCO procedure^46^ with slight modifications. Samples were dehydrated in increasing concentrations of methanol (20%, 40%, 60%, 80%, 100%) in H2O, and then washed in 100% methanol for 1 hr and incubated overnight in 66% dichloromethane (DCM) and 33% methanol at room temperature. The next day, samples were washed twice in 100% methanol for 30 min, cooled down to 4°C and bleached in chilled 5% H2O2 in methanol overnight at 4°C. Samples were rehydrated in decreasing concentrations of methanol (80%, 60%, 40%, 20%) in PBS with 0.2% Triton X-100, 1 hr each at room temperature, washed in PBS with 0.2% Triton X-100 for 2 × 1 hr at room temperature, followed by incubation in permeabilization solution (PBS with 0.2% Triton X-100, 20% Dimethyl sulfoxide, DMSO) and 0.3 M glycine over three nights at 37°C. This was followed by incubation in blocking solution (0.2% Triton X-100, 6% donkey serum, 10% DMSO and 0.02% sodium azide in PBS) over two nights at 37°C. Subsequently, samples were incubated with anti-GFP antibody (1:2000, gfp-1020, Aves Labs) in antibody buffer (2% Tween-20, 5% DMSO, 3% donkey serum, 0.01 mg heparin/ml in PBS) at 37°C for 7 days followed by a series of washes in washing solution (2% Tween-20 and 0.01 mg heparin/ml in PBS, 10 min, 20 min, 30 min, 1 hr, 2 hr and 2× overnight). Samples were incubated with secondary antibody anti-chicken-Cy5 (1:1000, 703-175-155, Jackson ImmunoResearch) in antibody buffer over seven nights at 37°C. This was followed by a series of washes in washing solution (10 min, 20 min, 30 min, 1 hr, 2 hr and three nights) and dehydration in increasing concentrations of methanol (20%, 40%, 60%, 80%, 100%) in H2O and incubation in 100% methanol overnight at room temperature. Samples were incubated in 66% DCM/33% methanol for 3 hr at room temperature followed by 100% DCM for 2 × 15 min. Finally, the samples were transferred to dibenzyl ether (DBE).

### Light sheet microscopy

Brains were imaged using a Bruker Luxendo LCS SPIM microscope. Samples were mounted with ventral side down in a cuvette and imaged in ethyl cinnamate (ECi). Horizontal images were acquired at 4.4x magnification in a z-stack at 10 μm intervals. Autofluorescence images were capturesd at 560 ± 20 nm (excitation) and 650 ± 25 nm (emission) wavelength and the GFP signal was imaged at 630 ± 15 nm excitation wavelength and 680 ± 15 nm emission wavelength.

### Atlas registration

Region delineation of whole-brain samples were obtained by atlas segmentation. Atlas annotations were obtained by alignment to a digital LSFM-based mouse brain atlas^47^. The atlas utilizes the common coordinate framework version 3 (CCFv3) developed by the Allen’s Institute of Brain Science^48^ and uses the same nomenclature. Registration between the atlas and the individual samples was performed by a global affine alignment followed by a local multi-resolution B-spline-based alignment, using the Elastix image registration toolbox^49^. The registration was based on the autofluorescence image volumes, which were pre-processed by down sampling to 20 μm isotropic resolution and contrast enhancement by contrast limited adaptive histogram equalization (CLAHE).

### Quantitative polymerase chain reaction (qPCR)

Tissues for qPCR analysis were collected into RNALater solution (Ambion) and stored at -20°C until processing. Total ribonucleic acid (RNA) was extracted using a Qiagen RNeasy Micro Kit (Qiagen) as per the manufacturer’s instructions in an RNase free environment. RNA quality and quantity was assessed spectrophotometrically using a NanoDrop ND-1000 (ThermoFisher Scientific) prior to the synthesis of single-stranded copy deoxyribonucleic acid (cDNA) using a High-Capacity cDNA Reverse Transcription Kit (Applied Biosystems) from 100-200 ng total RNA. qPCR was performed using SYBR Green chemistry on a QuantStudio 7.0 Flex Real-Time PCR System (Applied Biosystems) in 384 well plates. Relative gene expression was calculated using the 2^(-ΔΔCt) method using a *Gapdh* as a housekeeping gene as the expression of this gene did not change between groups. All primers were obtained from Sigma-Aldrich, sequences can be found in Table 1.

**Table 1.**
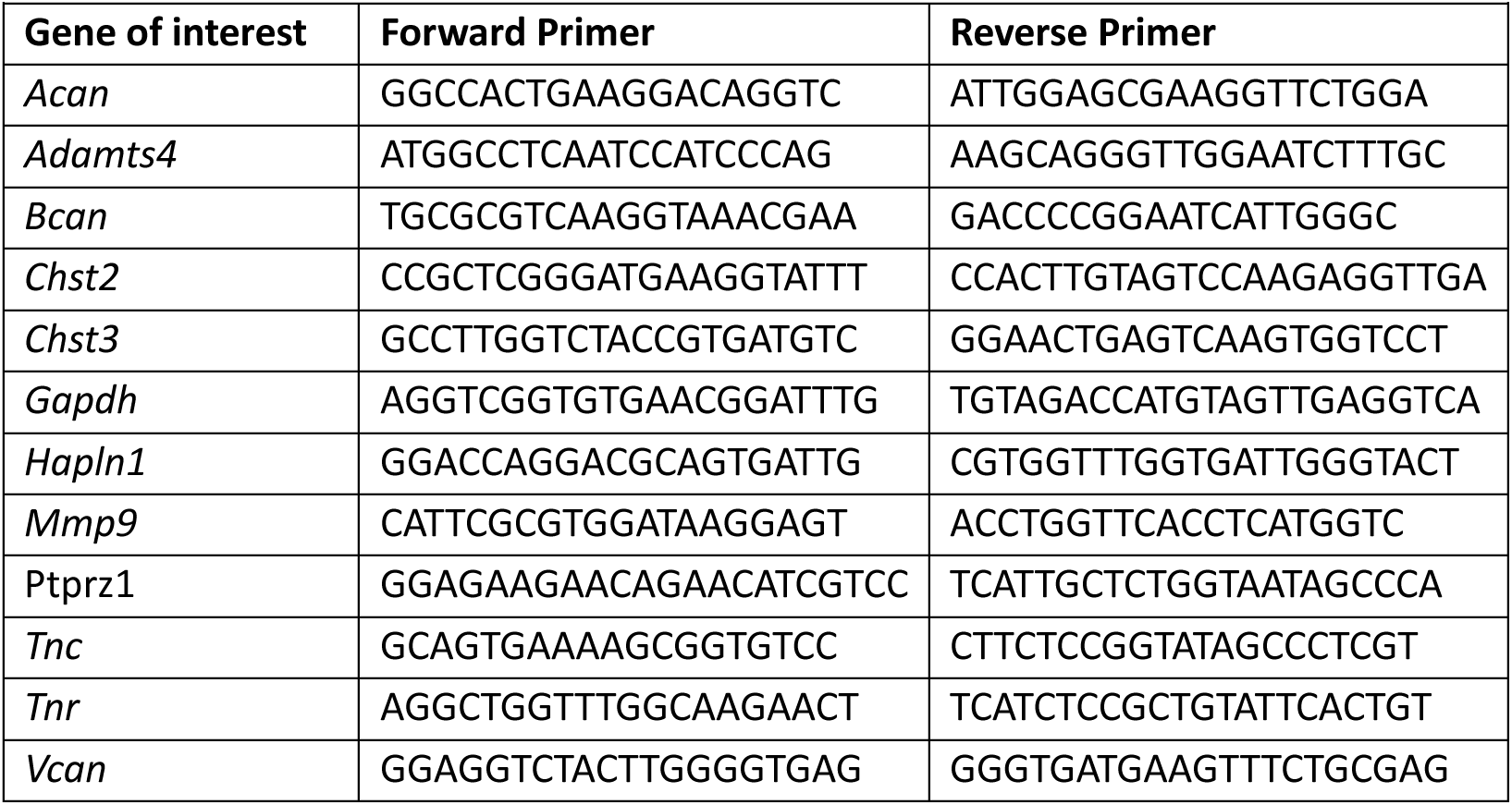
Primer sequences for SYBR Green qPCR.

### Analysis of HypoMap

Oligodendrocyte populations were isolated from HypoMap based on C7 cluster level labelling (C7-4 cluster is labelled as ‘Oligodendrocyte and precursor’. Visualisation of Adamts4 and other oligodendrocyte associated genes (Mog, Mbp, Bcas1, Bmp4, Tcf7l2, Pdgfra) was performed using Seurat v4 [https://doi.org/10.1016/j.cell.2021.04.048] and ggplot2. Average log-normalised expression of genes in each cluster at C66 level was used to create heatmaps of expression.

## Statistical analysis

All data visualisation and statistical analysis was performed in Prism 10 Software (GraphPad). Details of statistical tests are found in figure legends. All data sets were tested for normality and equality of variance prior to significance testing. Normality testing was performed using a Shapiro-Wilk test. Equality of variance was tested using Bartlett’s test or Brown-Forsythe test. Values ±2 standard deviations from the group mean were excluded from analysis, alternatively outliers were identified in GraphPad Prism using Grubb’s or ROUT’s test. All data are presented as mean ± standard error of the mean, n refers to the number of animals per group.

## Acknowledgements

We thank Professor Michael Schwartz at the University of Washington and Dr. Stavros Vagionitis, Mert Yucel and Professor Thora Karadottir at the Cambridge Stem Cell Institute for their insightful discussions and technical advice that have been essential to this project. We thank members of the Histopathology, Imaging and Disease Model Cores and the Core Biochemistry Assay Laboratory at the Institute of Metabolic Science Metabolic Research Laboratories for their assistance with this work. We also thank Julia Jones, Heather Zecchini and Huw Naylor at the Histopathology and ISH Core Facility and Light Microscopy Core Facility at the Cancer Research UK Cambridge Institute for their assistance with sm-FISH studies. We thank Louis Elfari at the Cambridge Stem Cell Institute Advanced Imaging Facility for processing and imaging SEM samples.

## Funding

This work was supported by a Medical Research Council grant (MR/S011552/1; CB), a Wellcome Trust PhD Studentship (108926/Z/15/Z; SB), Diabetes UK (22/0006401; CB), Medical Research Council Metabolic Disease Unit and Mouse Biochemistry Laboratory Grants (MC_UU_00014/5) and (MRC_MC_UU_12012/5), a Wellcome Trust Strategic Award (208363/Z/17/Z), a Wellcome Trust Investigator Award (108726/Z/15/Z; WDR), a Wellcome Trust Investigator Award (214286/Z/18/Z; HL, WDR) and a Sanming Award (SZSM201911003; WDR, HL) from the Municipal Government of Shenzhen, China, a Grant-in-Aid for Scientific Research (A) from the Japan Society for the Promotion of Science (JSPS; 20H00548; SH) and a Grant-in-Aid for Early-Career Scientists from the JSPS (23K16796; IS). EOS, CR, JS and MLE received funding from the Innovative Medicines Initiative 2 Joint Undertaking (JU) under grant agreement No. 777460 (HypoRESOLVE). The JU receives support from the European Union’s Horizon 2020 research and innovation programme and EFPIA and T1D Exchange, JDRF, International Diabetes Federation (IDF), The Leona M. and Harry B. Helmsley Charitable Trust. The University of Cambridge has received salary support for MLE through the National Health Service in the East of England through the Clinical Academic Reserve. The manuscript reflects the authors’ views, and the funders are not responsible for any use that may be made of the information it contains. For the purposes of open access, the authors have applied a CC-BY public copyright license to any Author Accepted Manuscript version arising from this submission.

## Author contributions

SB, MLE and CB designed experiments with input from HL and WDR. SB, EOS, CR, AT, JS, KI, GO, IS, SH, SS, SGN SN, MRV, JH, GKCD, BYHL, and CB performed and/or analysed experiments. SB, IS, SH, GSHY, MLE, HL, WDR and CB acquired funding. SB and CB wrote the original draft. All authors reviewed and edited the manuscript.

## Declaration of interests

The authors declare no competing interests.

## Data availability

Data available upon request to the corresponding author.

---

**Extended Data Figure 1:**
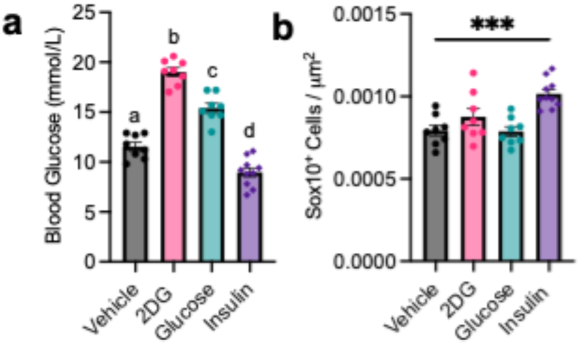
Glycaemia regulates median eminence oligodendrocyte lineage progression. (**a**) Blood glucose levels in C57BL/6J mice one hour after the intraperitoneal administration of vehicle (saline; 10 ml/kg), 2-deoxy-d-glucose (2DG; 250 mg/kg), glucose (2 g/kg) or insulin (0.75 U/kg). (**b**) Quantification of OL lineage cells (SOX10^­­^) in the ME one hour after vehicle, 2DG, glucose or insulin administration. All data presented as mean ± S.E.M.. Data analysed using one-way ANOVA with Tukey’s or Dunnett’s multiple comparisons test, ***p<0.001, n=8-10/group.

**Extended Data Figure 2:**
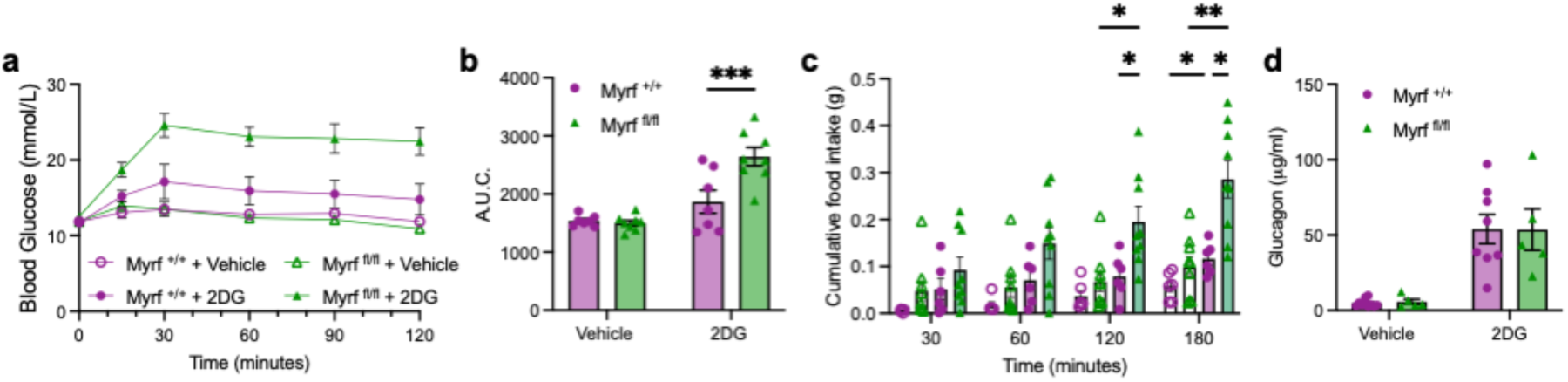
Adult oligodendrocyte differentiation is required for glucose homeostasis. **(a)** Blood glucose in *Myrf ^­­^*and *Myrf ^­­^*mice over 120 minutes after an intraperitoneal injection of vehicle (saline; 10 ml/kg) or 2-deoxy-d-glucose (2DG; 250 mg/kg) and (**b**) the resulting area under the curve. (**c**) Food intake over 180 minutes after vehicle or 2DG administration in *Myrf ^­­^*and *Myrf ^­­^*mice. (**d**) Plasma glucagon levels 30 minutes after vehicle or 2DG administration in *Myrf ^­­^*and *Myrf ^­­^*mice. Data presented as mean ± S.E.M.. Data analysed by unpaired student’s t test or two-way ANOVA with Tukey’s or Sidak’s multiple comparisons test.

**Extended Data Figure 3:**
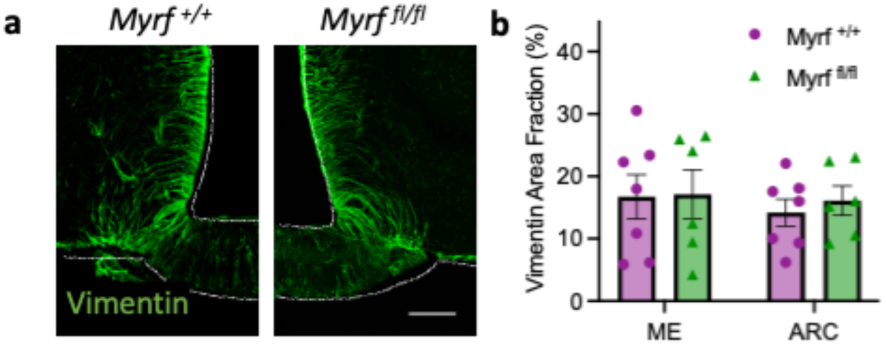
Adult oligodendrocyte differentiation regulates ARC glucose sensing mechanisms. Representative images of (**a**) Vimentin immunolabelling in the mediobasal hypothalamus of myelin regulatory factor (MYRF) conditional knockout mice (*Myrf ^­­^*) and controls (*Myrf ^­­^*) and (**b**) associated quantification. Scale bars represent 100 μm. Data presented as mean ± S.E.M. and analysed by unpaired students t-test, n=6-7/group.

**Extended Data Figure 4:**
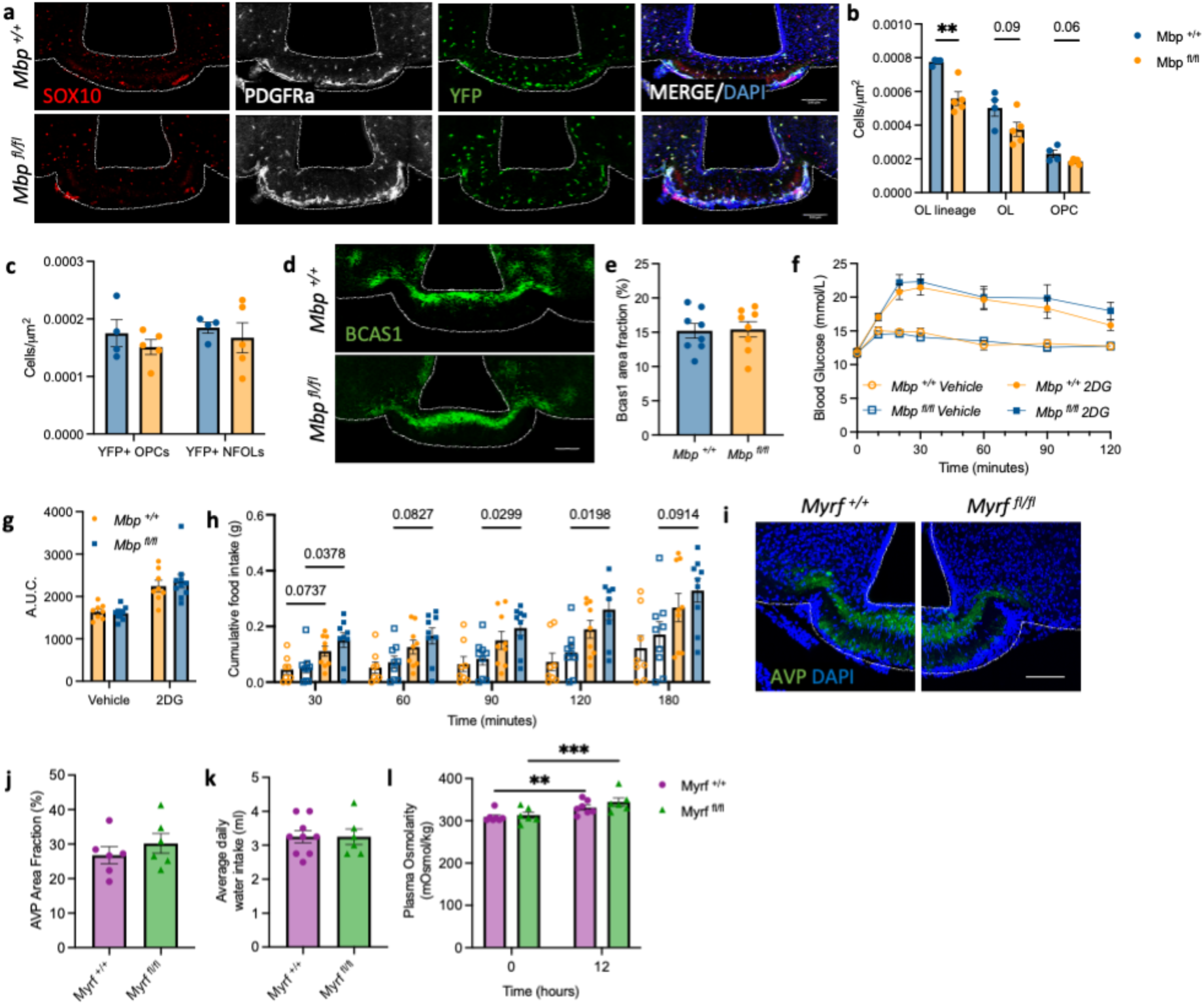
Adult myelin plasticity does not regulate glucose homeostasis. Immunolabelling against (**a**) pan-OL marker SOX10, OPC marker PDGFRa and yellow fluorescent protein (YFP) and (**d**) Breast carcinoma amplified sequence 1 (BCAS1) in the ME of MBP conditional knockout mice (*Mbp ^­­^*) and controls (*Mbp ^­­^*) and associated quantifications of (**b**) OL lineage cells, (**c**) OL lineage cells expressing YFP; NFOLs = newly formed OLs (SOX10+/PDGFRα-/YFP+) and (**e**) BCAS1 area fraction. (**f**) Blood glucose in *Mbp ^­­^*and *Mbp ^­­^*mice over 120 minutes after an intraperitoneal injection of vehicle (saline; 10 ml/kg) or 2-deoxy-d-glucose (2DG; 250 mg/kg) and (**g**) the resulting area under the curve. (**h**) Food intake over 180 minutes after vehicle or 2DG administration in *Mbp ^­­^*and *Mbp ^­­^*mice. (**i**) Representative images of arginine vasopressin immunolabelling in the mediobasal hypothalamus of myelin regulatory factor (MYRF) conditional knockout mice (*Myrf ^­­^*) and controls (*Myrf ^­­^*) and (**j**) associated quantification. (**k**) Daily water intake in *Myrf ^­­^*and *Myrf ^­­^*mice. (**l**) Plasma osmolarity in *Myrf ^­­^*and *Myrf ^­­^*mice at baseline and after 12 hours water deprivation. Data presented as mean ± S.E.M., scale bars represent 100 μm. Data analysed by unpaired student’s t test or two-way ANOVA with Tukey’s or Sidak’s multiple comparisons test, **p<0.01, n=4-9/group.

**Extended Data Figure 5:**
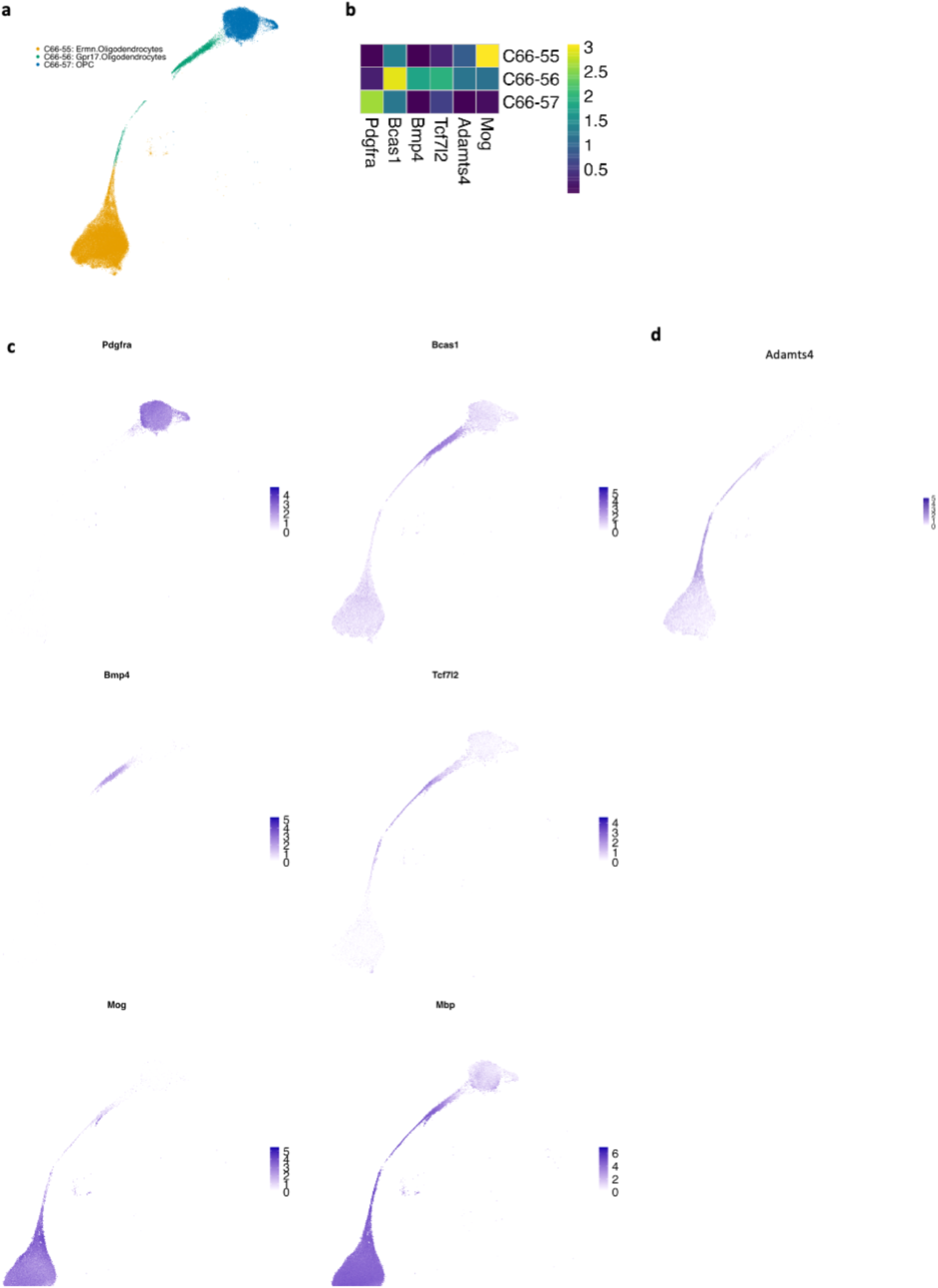
Oligodendrocyte progenitor cell differentiation regulates perineuronal net density and composition in the mediobasal hypothalamus. (a, b) Oligodendrocyte lineage population from HypoMap^­­^. Colours indicated C66 clustering, and highlight 3 distinct OL populations, representative of OPC, NFO and mature OL. (c) Heatmap displaying average log-normalized expression of marker genes of different stages of oligodendrocyte maturation in the 3 C66 OL clusters. (d) Log-normalized expression of Adamts4 in the OL population, demonstrating expression in NFO and mature OL populations.

**Extended Data Figure 6:**
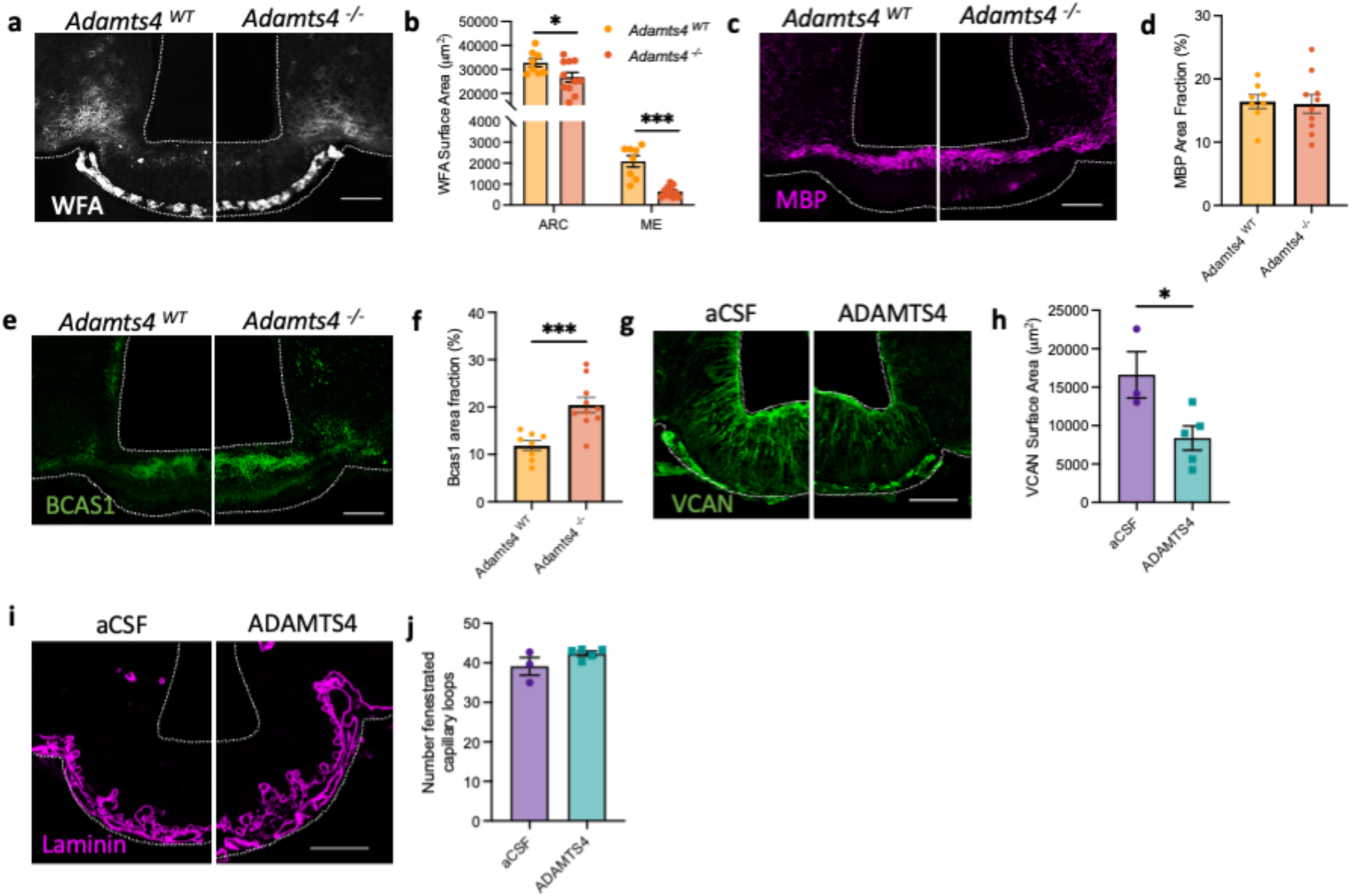
ADAMTS4 regulates the extracellular matrix and vascular permeability in the mediobasal hypothalamus. (**a**) Representative images of *Wisteria floribunda* agglutinin (WFA) lectin in the mediobasal hypothalamus (MBH) of ADAMTS4 knockout (*Adamts4^­­^*) mice compared to controls (*Adamts4^­­^*) and (**b**) associated quantification. Representative images of (**c**) MBP and (**e**) BCAS1 immunolabelling in the ME of *Adamts4^­­^*and *Adamts4^­­^*mice and (**d**, **f**) associated quantifications. Representative images of (**g**) VCAN and (**i**) laminin immunolabelling in the MBH of animals injected with aCSF or ADAMTS4 1 hour prior to sacrifice and (**h, j**) associated quantifications. Data presented as mean ± S.E.M. and analysed by unpaired student’s t-test or Mann-Whitney test, *p<0.05, ***p<0.001, ****p<0.0001, n=8-18/group. Scale bars represent 100 μm.

**Extended Data Figure 7:**
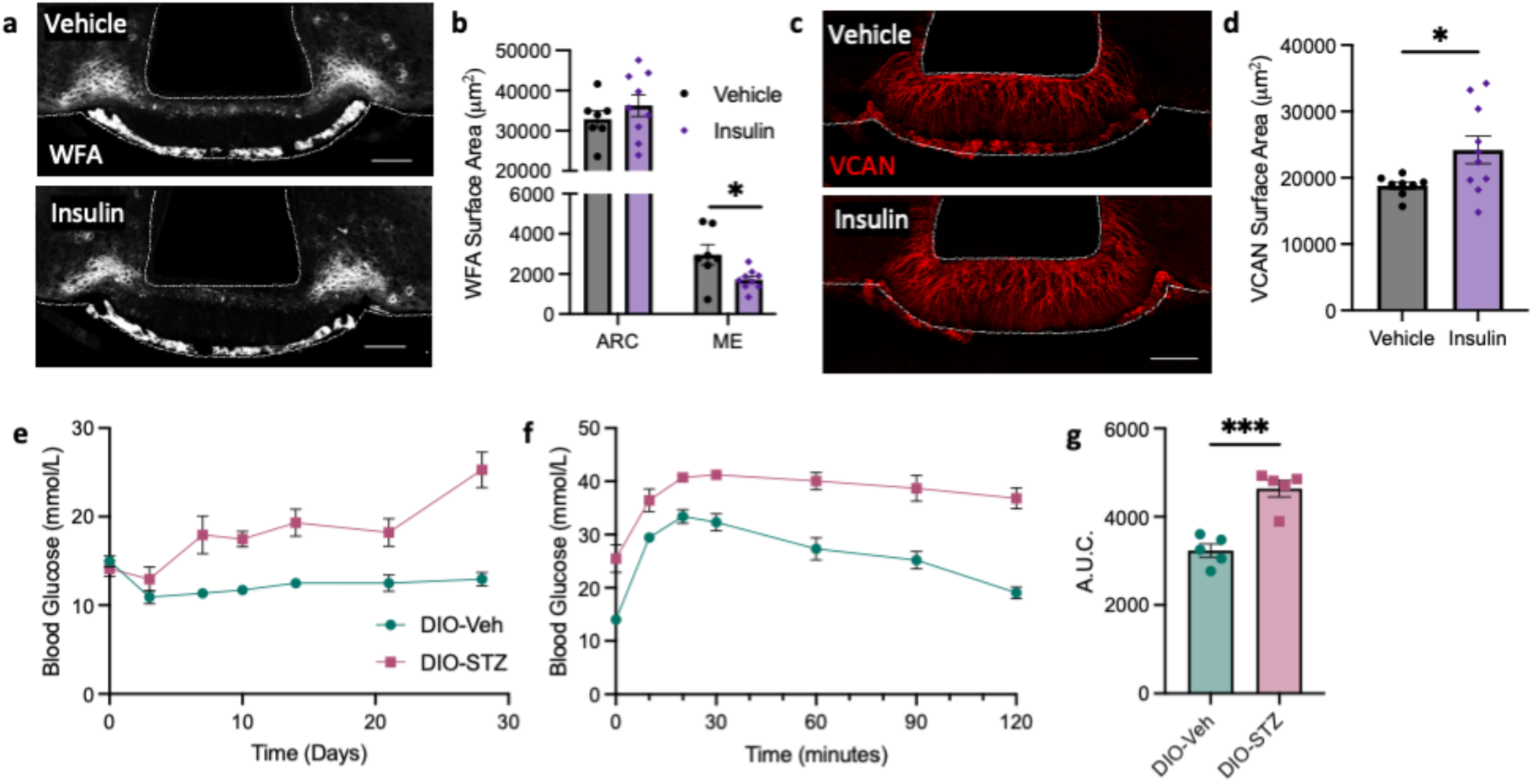
Median eminence ADAMTS4 expression is regulated by glycaemia. Representative images of (**a**) Wisteria floribunda agglutinin (WFA) lectin and (**c**) versican (VCAN) in the mediobasal hypothalamus (MBH) one hour after intraperitoneal (ip) administration of vehicle (saline; 10 ml/kg) or insulin (0.75 U/kg) to C57BL/6J mice and associated quantifications (**b**, **d**). (**e**) *Ad libitum* fed blood glucose levels in diet-induced obese (DIO) C57BL/6J mice over 4 weeks following ip administration of vehicle (saline, 10 ml/kg; DIO-STZ) or streptozotocin (STZ; 100 mg/kg; DIO-STZ). (**f**) Blood glucose levels in DIO-Veh and DIO-STZ mice over 2 hours following ip administration of glucose (1 g/kg) and (**g**) associated area under the curve. All data presented as mean ± S.E.M., scale bars represent 100 μm. Data analysed by unpaired student’s or Welch’s t-test or repeated measures 2-way ANOVA, *p<0.05, ***p<0.001, n=5-10/group.

## Notes

### Competing Interest Statement

The authors have declared no competing interest.

