## supplementary figures for "Adult oligodendrogenesis gates arcuate neuronal glucose sensing through remodelling of the blood-hypothalamus barrier via ADAMTS4"

Supplementary Materials

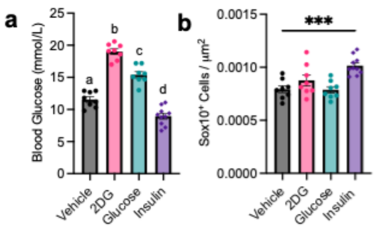


**Extended Data Figure 1: Glycaemia regulates median eminence oligodendrocyte lineage progression**

(**a**) Blood glucose levels in C57BL/6J mice one hour after the intraperitoneal administration of vehicle (saline; 10 ml/kg), 2-deoxy-d-glucose (2DG; 250 mg/kg), glucose (2 g/kg) or insulin (0.75 U/kg). (**b**) Quantification of OL lineage cells ($Sox10^+$) in the ME one hour after vehicle, 2DG, glucose or insulin administration. All data presented as mean ± S.E.M.. Data analysed using one-way ANOVA with Tukey's or Dunnett's multiple comparisons test, ***$p<0.001$, n=8-10/group.

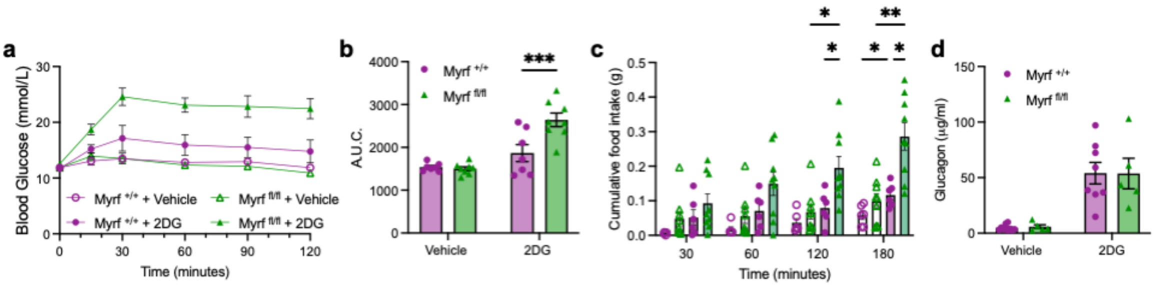


**Extended Data Figure 2: Adult oligodendrocyte differentiation is required for glucose homeostasis**

(**a**) Blood glucose in *Myrf* $^{+/+}$ and *Myrf* $^{fl/fl}$ mice over 120 minutes after an intraperitoneal injection of vehicle (saline; 10 ml/kg) or 2-deoxy-d-glucose (2DG; 250 mg/kg) and (**b**) the resulting area under the curve. (**c**) Food intake over 180 minutes after vehicle or 2DG administration in *Myrf* $^{+/+}$ and *Myrf* $^{fl/fl}$ mice. (**d**) Plasma glucagon levels 30 minutes after vehicle or 2DG administration in *Myrf* $^{+/+}$ and *Myrf* $^{fl/fl}$ mice. Data presented as mean ± S.E.M.. Data analysed by unpaired student's t test or two-way ANOVA with Tukey's or Sidak's multiple comparisons test.

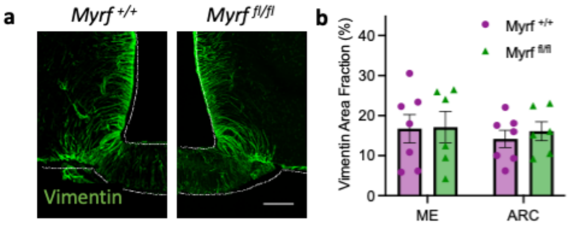


**Extended Data Figure 3: Adult oligodendrocyte differentiation regulates ARC glucose sensing mechanisms**

Representative images of (**a**) Vimentin immunolabelling in the mediobasal hypothalamus of myelin regulatory factor (MYRF) conditional knockout mice (*Myrf* $^{fl/fl}$) and controls (*Myrf* $^{+/+}$) and (**b**) associated quantification. Scale bars represent 100 µm. Data presented as mean ± S.E.M. and analysed by unpaired students t-test, n=6-7/group.

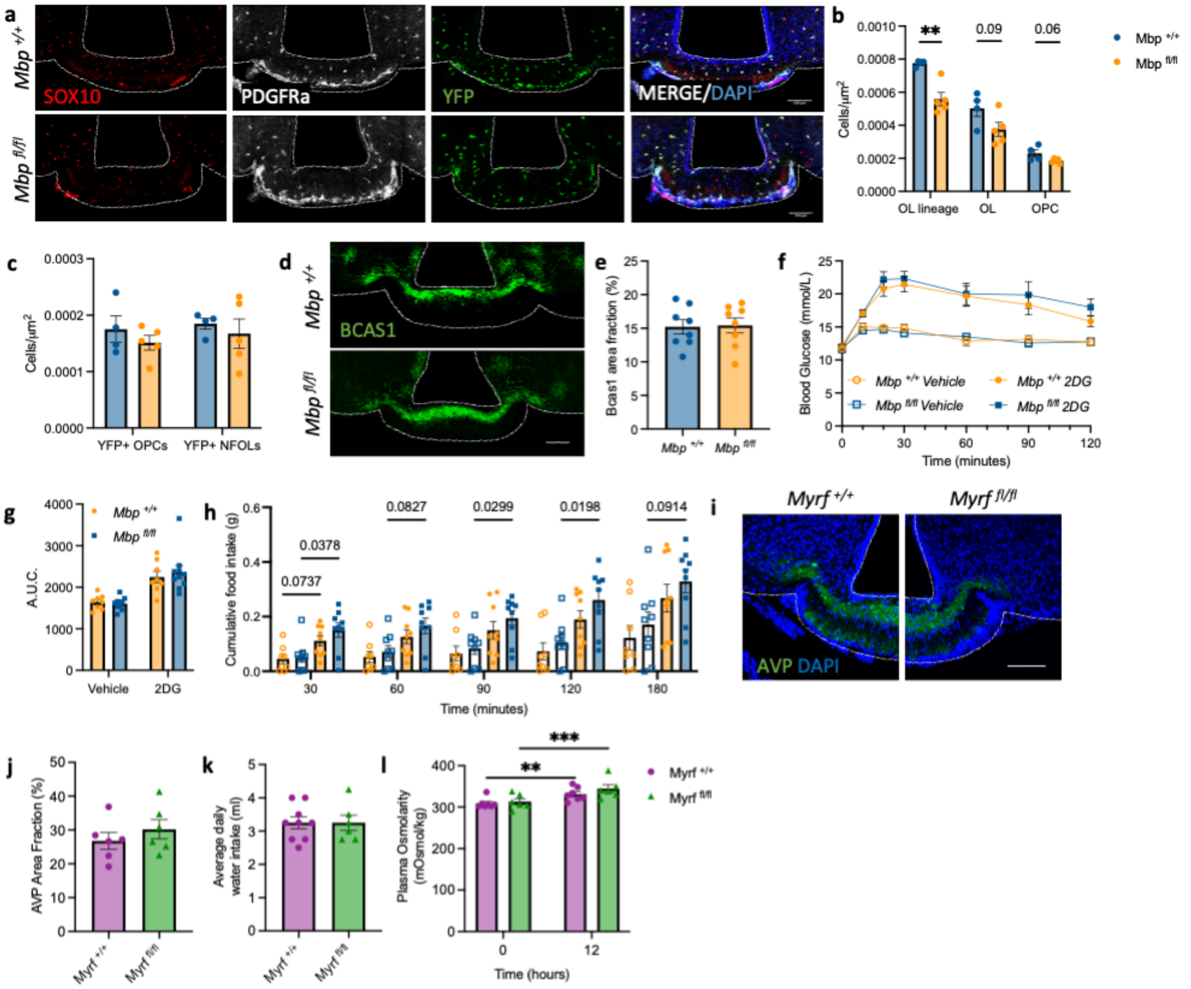


**Extended Data Figure 4: Adult myelin plasticity does not regulate glucose homeostasis**

Immunolabelling against (**a**) pan-oligodendrocyte (OL) marker, SRY-box transcription factor 10 (SOX10), OL progenitor cell (OPC) marker, platelet derived growth factor receptor alpha (PDGFRa) and yellow fluorescent protein (YFP) and (**d**) Breast carcinoma amplified sequence 1 (BCAS1) in the ME of MBP conditional knockout mice (*Mbp* $^{fl/fl}$) and controls (*Mbp* $^{+/+}$) and associated quantifications of (**b**) OL lineage cells, (**c**) OL lineage cells expressing YFP; NFOLs = newly formed OLs (SOX10+/PDGFRα-/YFP+) and (**e**) BCAS1 area fraction. (**f**) Blood glucose in *Mbp* $^{+/+}$ and *Mbp* $^{fl/fl}$ mice over 120 minutes after an intraperitoneal injection of vehicle (saline; 10 ml/kg) or 2-deoxy-d-glucose (2DG; 250 mg/kg) and (**g**) the resulting area under the curve. (**h**) Food intake over 180 minutes after vehicle or 2DG administration in *Mbp* $^{+/+}$ and *Mbp* $^{fl/fl}$ mice. (**i**) Representative images of arginine vasopressin immunolabelling in the mediobasal hypothalamus of myelin regulatory factor (MYRF) conditional knockout mice (*Myrf* $^{fl/fl}$) and controls (*Myrf* $^{+/+}$) and (**j**) associated quantification. (**k**) Daily water intake in *Myrf* $^{fl/fl}$ and *Myrf* $^{+/+}$ mice. (**l**) Plasma osmolarity in *Myrf* $^{fl/fl}$ and *Myrf* $^{+/+}$ mice at baseline and after 12 hours water deprivation. Data presented as mean ± S.E.M., scale bars represent 100 µm. Data analysed by unpaired student's t test or two-way ANOVA with Tukey's or Sidak's multiple comparisons test, **p<0.01, n=4-9/group.

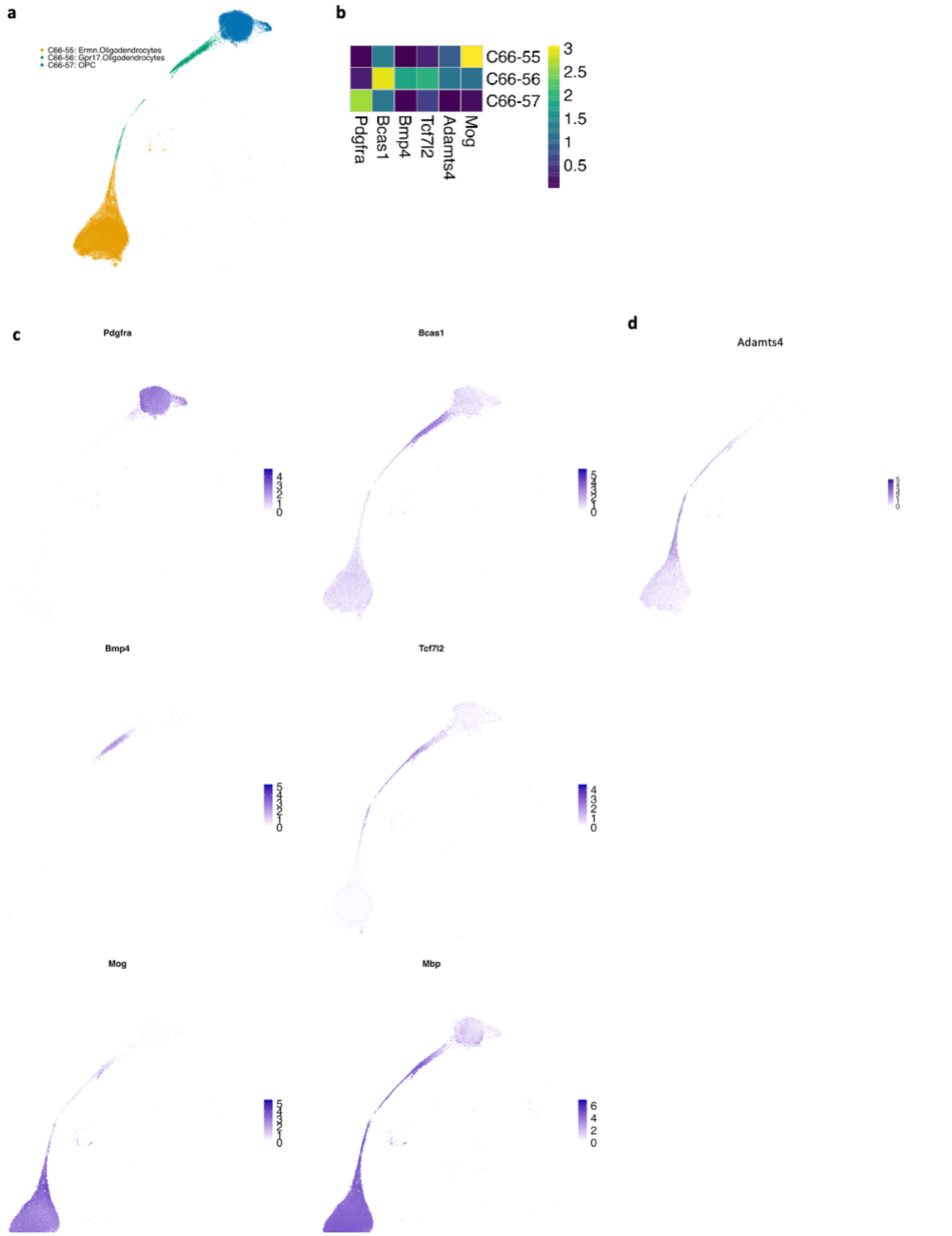


**Extended Data Figure 5: Oligodendrocyte progenitor cell differentiation regulates perineuronal net density and composition in the mediobasal hypothalamus.** **(a, b)** Oligodendrocyte lineage population from HypoMap[19]. Colours indicated C66 clustering, and highlight 3 distinct OL populations, representative of OPC, NFO and mature OL. (**c)** Heatmap displaying average log-normalized expression of marker genes of different stages of oligodendrocyte maturation in the 3 C66 OL clusters. (**d)** Log-normalized expression of Adamts4 in the OL population, demonstrating expression in NFO and mature OL populations.

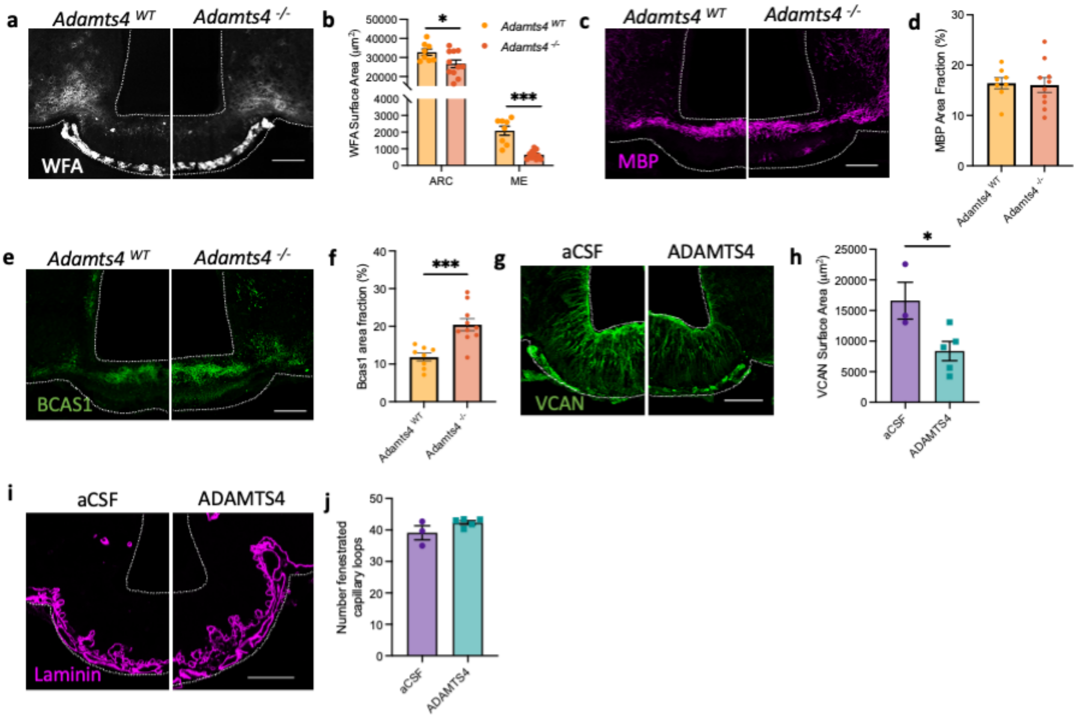


**Extended Data Figure 6: ADAMTS4 regulates the extracellular matrix and vascular permeability in the mediobasal hypothalamus**

(**a**) Representative images of *Wisteria floribunda* agglutinin (WFA) lectin in the mediobasal hypothalamus (MBH) of ADAMTS4 knockout (*Adamts4*$^{-/-}$) mice compared to controls (*Adamts4*$^{WT}$) and (**b**) associated quantification. Representative images of (**c**) MBP and (**e**) BCAS1 immunolabelling in the ME of *Adamts4*$^{-/-}$ and *Adamts4*$^{WT}$ mice and (**d**, **f**) associated quantifications. Representative images of (**g**) VCAN and (**i**) laminin immunolabelling in the MBH of animals injected with aCSF or ADAMTS4 1 hour prior to sacrifice and (**h, j**) associated quantifications. Data presented as mean ± S.E.M. and analysed by unpaired student's t-test or Mann-Whitney test, *p<0.05, ***p<0.001, ****p<0.0001, n=8-18/group. Scale bars represent 100 µm.

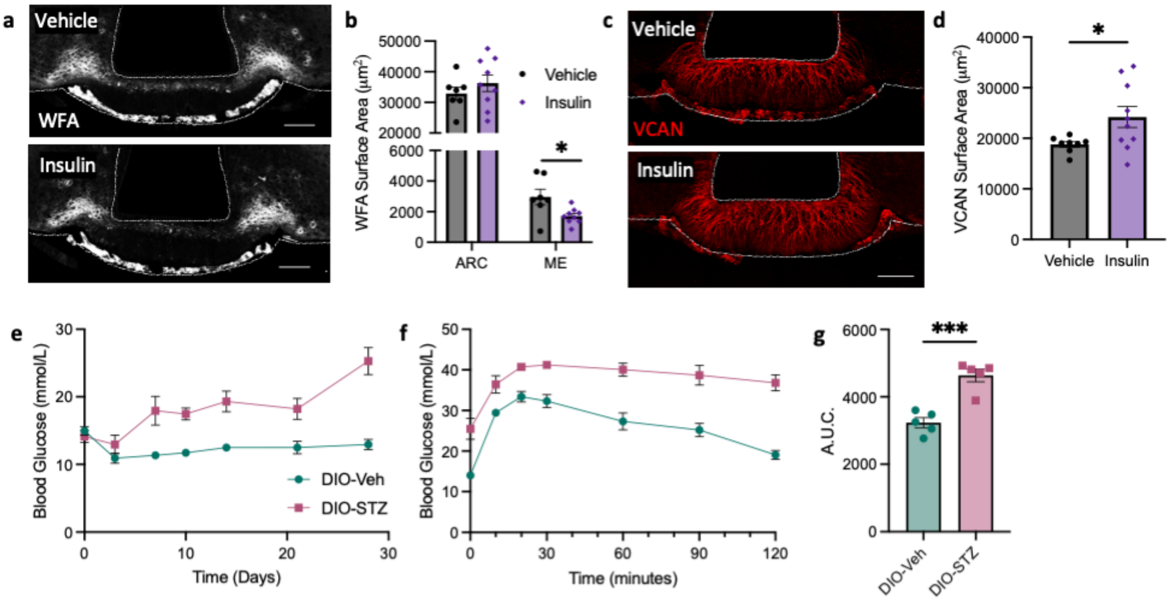


**Extended Data Figure 7: Median eminence ADAMTS4 expression is regulated by glycaemia**

Representative images of (**a**) Wisteria floribunda agglutinin (WFA) lectin and (**c**) versican (VCAN) in the mediobasal hypothalamus (MBH) one hour after intraperitoneal (ip) administration of vehicle (saline; 10 ml/kg) or insulin (0.75 U/kg) to C57BL/6J mice and associated quantifications (**b**, **d**). (**e**) *Ad libitum* fed blood glucose levels in diet-induced obese (DIO) C57BL/6J mice over 4 weeks following ip administration of vehicle (saline, 10 ml/kg; DIO-STZ) or streptozotocin (STZ; 100 mg/kg; DIO-STZ). (**f**) Blood glucose levels in DIO-Veh and DIO-STZ mice over 2 hours following ip administration of glucose (1 g/kg) and (**g**) associated area under the curve. All data presented as mean ± S.E.M., scale bars represent 100 µm. Data analysed by unpaired student's or Welch's t-test or repeated measures 2-way ANOVA, *p<0.05, ***p<0.001, n=5-10/group.

Supplementary video 1:

Distribution of mGFP fluorescent in a brain from *Pdgfrα-CreERT2;taumGFP* mice dosed with tamoxifen at P60 as previously described[10] , harvested at P90 , cleared using iDISCO and imaged using light-sheet 3D imaging.

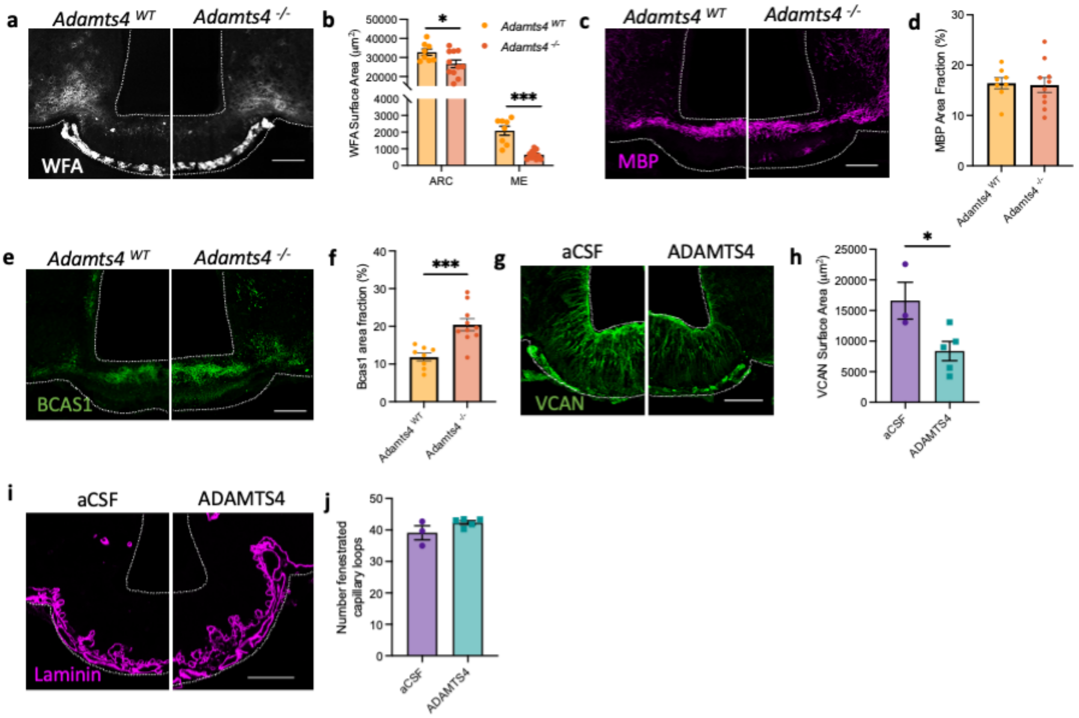


**Extended Data Figure 6: ADAMTS4 regulates the extracellular matrix and vascular permeability in the mediobasal hypothalamus**

(**a**) Representative images of *Wisteria floribunda* agglutinin (WFA) lectin in the mediobasal hypothalamus (MBH) of ADAMTS4 knockout (*Adamts4*$^{-/-}$) mice compared to controls (*Adamts4*$^{WT}$) and (**b**) associated quantification. Representative images of (**c**) MBP and (**e**) BCAS1 immunolabelling in the ME of *Adamts4*$^{-/-}$ and *Adamts4*$^{WT}$ mice and (**d**, **f**) associated quantifications. Representative images of (**g**) VCAN and (**i**) laminin immunolabelling in the MBH of animals injected with aCSF or ADAMTS4 1 hour prior to sacrifice and (**h, j**) associated quantifications. Data presented as mean ± S.E.M. and analysed by unpaired student's t-test or Mann-Whitney test, *p<0.05, ***p<0.001, ****p<0.0001, n=8-18/group. Scale bars represent 100 µm.

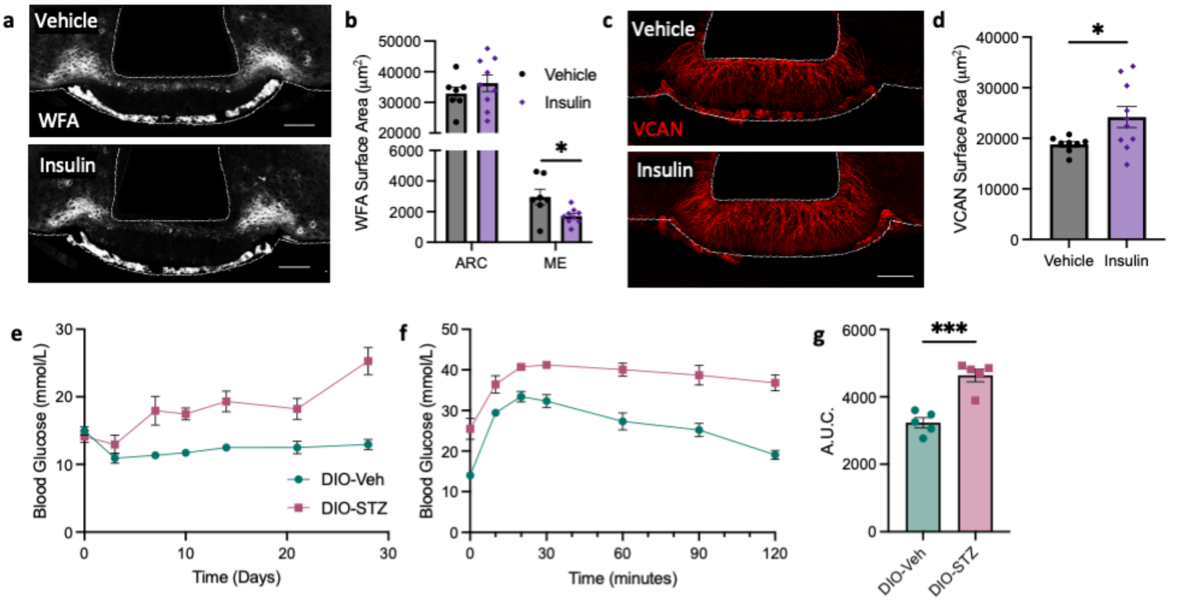


**Extended Data Figure 7: Median eminence ADAMTS4 expression is regulated by glycaemia**

Representative images of (**a**) Wisteria floribunda agglutinin (WFA) lectin and (**c**) versican (VCAN) in the mediobasal hypothalamus (MBH) one hour after intraperitoneal (ip) administration of vehicle (saline; 10 ml/kg) or insulin (0.75 U/kg) to C57BL/6J mice and associated quantifications (**b**, **d**). (**e**) *Ad libitum* fed blood glucose levels in diet-induced obese (DIO) C57BL/6J mice over 4 weeks following ip administration of vehicle (saline, 10 ml/kg; DIO-STZ) or streptozotocin (STZ; 100 mg/kg; DIO-STZ). (**f**) Blood glucose levels in DIO-Veh and DIO-STZ mice over 2 hours following ip administration of glucose (1 g/kg) and (**g**) associated area under the curve. All data presented as mean ± S.E.M., scale bars represent 100 µm. Data analysed by unpaired student's or Welch's t-test or repeated measures 2-way ANOVA, *p<0.05, ***p<0.001, n=5-10/group.
